# Integrated multi-omics reveals organ-specific jasmonate signaling in Arabidopsis under combined heat and phosphate starvation

**DOI:** 10.64898/2026.11.16.766386

**Authors:** Victoria Baca-González, Francesco Blasio, Mónica Lanza, Alberto Lozano-Enguita, Irene Torregrosa Gómez-Meana, Adrián Sabater, Sara Navarro-Neila, Jose M. Garcia-Mina, Angel M. Zamarreño, Maria Garnica-Ochoa, Elena Caro, Juan C. del Pozo

## Abstract

Phosphate (Pi) limitation is a major constraint on plant productivity and is aggravated by heat stress. In nature, soil buffers air temperature, creating a vertical thermal gradient that shields roots from extreme heat; thus, heat and Pi starvation, although often co-occurring, are perceived mainly by different organs, shoots and roots respectively. How plants integrate these organ-specific stresses into a coordinated response remains poorly understood. Here, we investigated how Arabidopsis responds to combined Pi deficiency and elevated temperature, using a root-zone temperature gradient and an integrated analysis of ion accumulation, phytohormone levels and transcriptomic profiles in roots and shoots. The combined stress elicited a distinct response that cannot be explained by the addition of individual stress effects. Jasmonate levels changed in an organ-specific manner, with JA-Ile accumulating in roots and JA declining in shoots, and mutants with reduced (*jar1-1*) or constitutive (*jazQ*) JA signaling showed altered transcriptional, growth and leaf-temperature responses. These results identify jasmonic acid (JA) as a modulator of the balance between acclimation and growth under combined nutritional and thermal limitation, advancing our understanding of how plants cope with the multi-stress scenarios driven by climate change.

**Short summary:** Heat and phosphate (Pi) starvation are sensed mainly by shoots and roots, respectively. Using a root-zone temperature gradient, we show that their combination triggers specific ionomic, hormonal and transcriptional responses and an organ-specific rewiring of jasmonates that modulates the balance between stress acclimation and growth.

## Introduction

Phosphorus is an essential nutrient for plant growth and development. Plants have evolved complex regulatory mechanisms to improve the uptake of inorganic phosphate (PO_4_^3-^; Pi) from the rhizosphere and its internal use (Baek et al., 2017). When Pi levels fall below a critical threshold, plants activate the phosphate starvation response (PSR), which mobilizes Pi from the rhizosphere and reallocates it from storage to actively growing tissues. The PSR involves metabolic readjustment, morphological changes and specific transcriptional programs, mainly regulated by the PHR1-miR399-PHO2 module (Bari et al., 2006; Bustos et al., 2010; Lopez-Bucio et al., 2003; Peret et al., 2011; Rubio et al., 2001; Secco and Whelan, 2014). Many of these morphological changes, such as increased lateral root (LR) density and root hair formation, enlarge the root surface to explore the soil for Pi. These morphological, metabolic and gene expression adjustments are influenced by root illumination, and the combination of Pi starvation and root illumination generates a specific plant response (Silva-Navas et al., 2019).

Climate change is increasing the frequency of high-temperature extremes, such as heat waves, which strongly reduce plant development and agricultural yield (Bhattacharya, 2019; Gray and Brady, 2016; Miller et al., 2021). Heat stress impairs protein synthesis and folding, alters enzyme activity and damages membranes and cellular structures through oxidation (Hasanuzzaman et al., 2013). Plants cope with these challenges through changes in gene expression, metabolism and morphology (Zhao et al., 2020). In natural and agricultural environments, Pi starvation rarely occurs in isolation. High temperatures can trigger the PSR by impairing Pi uptake by roots (Bechtaoui et al., 2021; Giri et al., 2017) and by disrupting Pi transport and distribution within the plant, reducing photosynthetic efficiency, energy production and growth (Bechtaoui *et al*., 2021; Mishra et al., 2023). Although Pi starvation and heat stress have been studied individually, how plants integrate responses to their combination remains poorly understood. In sheepgrass (*Leymus chinensis*), the combination of both stresses reduces shoot growth and biomass more than either stress alone (Li et al., 2018). In *Brassica napus*, the impact of combined drought and heatwave is strongly modulated by nutrient availability, particularly during recovery (Dikšaitytė et al., 2020), and in cool-season grasses heat-induced iron deficiency is a critical component of high-temperature stress (Minami et al., 2025).

Given the increasing frequency of extreme temperatures and the persistent low Pi availability of agricultural soils, dissecting plant responses to combined Pi starvation and heat is crucial for developing climate-resilient and nutrient-efficient crops. Here, we found that combined Pi deficiency and heat leads to a severe nutrient imbalance in roots and shoots and to marked changes in hormone levels. The transcriptional response to the stress combination differed markedly from those to the individual stresses, and jasmonic acid (JA) levels and signaling were altered by the combined stress. Transcriptomic profiling of WT and *jar1-1* plants, deficient in JA-Ile biosynthesis, showed that balanced JA-Ile signaling is required for a proper transcriptional response to Pi starvation, heat and their combination. In soil, *jar1-1* showed reduced growth, in contrast to its higher biomass in vitro, highlighting the relevance of experimental setups close to natural conditions. Together with the restricted growth and lower leaf temperatures under Pi-sufficient conditions of the *jazQ* mutant, which lacks five *JAZ* repressors and shows constitutive JA signaling, these results suggest that proper JA signaling is needed for the plant response to combined Pi deficiency and heat stress. Our results identify organ-specific modulation of jasmonates, with JA-Ile accumulating in roots and JA declining in shoots, as a mechanism by which plants integrate spatially separated nutrient and heat signals to balance stress acclimation and growth.

## Materials and Methods

### Plant material

*Arabidopsis thaliana* Columbia (Col-0) was used as wild type (WT). The *jar1-1* mutant (Staswick et al., 2002) was used for transcriptomic and phenotypic analyses, and the *jazQ* mutant (Campos et al., 2016) for phenotyping. Seeds were surface-sterilized with 70% commercial bleach for 10 min, rinsed three times with sterile water and sown on vertically oriented 12 × 12 cm plates containing half-strength Murashige and Skoog (½ MS) medium without phosphate, 0.05% MES, 1% sucrose and 1% agar (Difco, Sparks, MD, USA), pH 5.7. Filter-sterilized KH₂PO₄ was added to the autoclaved medium at the indicated concentrations, and potassium was compensated with equimolar KCl. Plants were grown in the D-Root device (Silva-Navas et al., 2015) to keep roots in darkness or, when indicated, in the TGRooZ (González-García et al., 2022) to generate a temperature gradient in the root zone. For temperature gradient experiments, seedlings were grown at 22°C in the TGRooZ (22G) in HPi or LPi for 4 days and then half of the plates were maintained at 22G (22G-HPi, or 22G-LPi) and half transferred to the 32°C TGRooZ (32G, in the corresponding Pi condition: 32G-HPi or 32G-LPi). The 32G gradient ranged from 32°C in the upper part to 24°C in the lower part of the plate, and the 22G gradient from 22°C to 17–18°C. In those experiments using homogeneous temperature in roots and shoots, seedlings were grown at 22°C (22SR) in D-Root devices, and half of the plates were transferred to 32°C (32SR). After identifying 30 µM as the threshold Pi concentration that activates the phosphate starvation response at 32G (Fig. S1), we set high Pi (HPi) and low Pi (LPi) conditions at 625 µM and 30 µM Pi, respectively (Lozano-Enguita et al., 2026). Three biological replicates per condition were used, except for hormone and element quantification (four replicates). Seedlings were grown under a 16-h light/8-h dark photoperiod with a light fluence rate of 120 µmol m-2 s-1 at the bottom of the plate, provided by LED tubes.

### Root growth assays and microscopic analysis

Primary root and total LR length were measured with ImageJ. Emerged lateral roots were counted under a stereomicroscope.

### Plant phenotyping

WT (Col-0), *jar1-1* and *jazQ* seedlings were germinated on ½ MS medium with HPi or LPi and transferred to soil 7 days after germination. The soil substrate consisted of blond peat with low nutrient content (Turbaflor™; Table S1) mixed 5:1 (v/v) with clean, sterilized river sand. To generate high- and low-Pi conditions, the substrate was fertilized with either 2.5 g/L of NPK fertilizer (high-Pi treatment; 16% N: 7% KNO3 + 9% NH4NO3; 9% P2O5; 12% K2O; 2% MgO; 0.02% B; 0.005% Cu; 0.45% Fe; 0.06% Mn; 0.02% Mo; 0.015% Zn) or 2.3 g/L of an equivalent NK fertilizer without phosphorus (low-Pi treatment). Plants were transplanted into 0.5-L pots, watered to field capacity and maintained at 24°C for 2 days; one third of the plants were then kept at 24°C (24SR) and the other two thirds were shifted to homogeneous 34°C (34SR) or to 34°C in the TGRooZ (34G) for a further 11 days (n = 13 plants per genotype and condition). Rosette growth was monitored with an RGB camera. Individual plants were segmented to obtain total rosette area, and leaves were counted by applying a watershed algorithm to the binary mask, using LemnaTec software. The thermal images were acquired with FLIR96-24 camera at 1 pm (5 h after the start of the light period) and analyzed with the FLIR suite software.

### Hormone profiling

Indole-3-acetic acid (IAA), indole-3-butyric acid (IBA), abscisic acid (ABA), jasmonic acid (JA), jasmonoyl-L-isoleucine (JA-Ile), salicylic acid (SA), trans-zeatin (tZ), trans-zeatin riboside (tZR), cis-zeatin (cZ), cis-zeatin riboside (cZR), N6-isopentenyladenine (iP) and isopentenyladenosine (iPR) were quantified separately in roots and shoots of 22G-HPi (control), 22G-LPi, 32G-HPi and 32G-LPi seedlings as described (Rubio et al., 2026).

### Free phosphate quantification and element quantification

Inorganic phosphate was quantified as described (Ames, 1966) in seedlings grown for 4 days on ½ MS medium with the indicated Pi concentrations in the D-Root device and then transferred to the indicated temperature regimes for 6 days. Free Pi was expressed per mg of fresh weight and, owing to variability between experiments, also relative to the control as indicated.

Ionomic analyses were performed as described (Silva-Navas *et al*., 2015). WT seedlings were grown with 625 µM or 30 µM Pi in D-Root devices for 4 days and transferred to 22G or 32G for 6 days. Roots and shoots were collected separately, dried at 100 °C for 24–48 h in an air oven and digested with 3 mL of HNO3 (Sigma-Aldrich Trace Metal grade) at 100 °C in a thermostatized bath for 4 h. Samples were diluted to 10 mL with 18 MΩ water and analyzed by Inductively Coupled Plasma-Optical Emission Spectrometry (ICP-OES) on a Thermo Scientific iCAP 6500 at the following wavelengths: B, 208.9 nm; Ca, 184.0 and 317.9 nm; Fe, 240.4 and 259.9 nm; K, 766.4 nm; Mg, 279.0 and 285.2 nm; Mn, 293.9 nm; Mo, 203.8 nm; Na, 589.5 nm; P, 177.4 and 178.2 nm; S, 182.0 nm; Zn, 206.2 nm; and Si, 251.6 nm.

### Differential gene expression analyses

Roots and shoots of WT or *jar1-1* seedlings grown for 4 days at 22SR in HPi or LPi and then transferred to 22G or 32G for 6 days (22G-HPi, 32G-HPi, 22G-LPi and 32G-LPi; three biological replicates per condition) were separated with sterile scalpels, frozen in liquid nitrogen and stored at −80°C. Total RNA was extracted using the RNeasy Mini Kit (Qiagen, cat. no. 74104), and approximately 20 μg was sent to Beijing Genomics Institute (BGI-Shenzhen) for library construction and sequencing. Poly-A+ RNA was isolated by oligo-dT chromatography, fragmented and converted to double-stranded cDNA using random hexamer primers, followed by end repair, 3′ end adenylation and adapter ligation. cDNA fragments were selected by agarose gel extraction, enriched by PCR and sequenced on an Illumina platform (150-bp paired-end reads; ∼30 M reads per sample; average quality 96%). These data were generated with the same experimental design as in a previous study (Lozano-Enguita *et al*., 2026), and sequencing data are deposited in GEO under accession number GSE327853. For the comparison between WT (Col-0) and *jar1-1*, a new set of WT and mutant seedlings was grown and processed as described above.

Read quality before and after trimming was assessed with FastQC v0.12.1. Reads were aligned to the Arabidopsis thaliana TAIR10 genome using HISAT2 v2.2.1 with default parameters, and SAM files were converted to BAM format, coordinate-sorted and indexed with SAMtools v1.21. Reads were assigned to genes using featureCounts from the Subread v2.0.6 package (options -p -t exon -g gene_id) with the Araport11 annotation GTF file.

Differential gene expression analysis was performed using DESeq2 v1.50.2 in R/Bioconductor with a factorial design including main effects and interaction terms for tissue type, temperature and phosphate condition. Genes with at least 10 reads in at least three samples were retained. Normalization and dispersion estimation used default parameters, with 22G and HPi as control reference levels within each tissue. Genes with adjusted p-values <0.05 and |log2 fold change| > 0.75 (∼1.7-fold) were considered differentially expressed. Principal component analysis (PCA) of variance-stabilized transformed (VST) data was used to assess sample clustering and potential batch effects.

### Gene Ontology and Statistical Analyses

GO enrichment analyses were performed with clusterProfiler (REF) using the Arabidopsis genome as background; p-values were adjusted by the Benjamini–Hochberg method, and terms with adjusted p < 0.05 were considered enriched. Heatmaps were generated using the One-for-All tool (Guit1234). Statistical analyses were performed using GraphPad Prism 8 (GraphPad Software, San Diego, CA, USA). Two groups were compared with Student’s t-test, and multiple groups by one-way/two-way analysis of variance (ANOVA) followed by Tukey’s or Dunnett’s post hoc test (P < 0.05). Ionomic and hormone data were log10-transformed and analyzed by one-way ANOVA followed by Dunnett’s test versus the control (22G-HPi). Daily root growth (Fig. S2A) was analyzed for each day by two-way ANOVA (temperature × Pi) and Tukey’s HSD test among the eight conditions using statsmodels (v0.14.1) in Python. Rosette fresh weight in soil was analyzed by three-way ANOVA (genotype × temperature × Pi). Different letters indicate means that differ significantly.

## Results

### A temperature gradient within the root growth zone modulates root and shoot responses to combined phosphate starvation and heat stress

Because heat stress disrupts nutrient homeostasis, we examined the developmental, physiological and molecular responses of *Arabidopsis thaliana* seedlings to combined Pi deficiency and elevated shoot temperature. To mimic soil conditions, we used the D-Root and TGRooZ devices, which keep roots in darkness and generate a temperature gradient along the root system. Shoots were exposed to 22°C or 32°C while roots grew in gradients from 22 to 17–18°C (22G) or from 32 to 24°C (32G), respectively. For comparison, plants were also grown with shoots and roots at a uniform 22°C or 32°C (22SR and 32SR), although these conditions do not reflect a natural soil environment. Using the Pi starvation-inducible *pSPX3:GUS* reporter line (Duan et al., 2008), we found that 30 μM Pi (hereafter low phosphate, LPi) was the threshold concentration that triggered the PSR under 32G conditions (Fig. S1). Root growth and lateral root (LR) formation were strongly reduced only under homogeneous 32SR conditions (Fig. 1A,B), whereas the TGRooZ regimes had minimal impact. Although 22SR and 32G seedlings showed more LRs irrespective of Pi availability, lateral root density (LRD) did not change among conditions (Fig. 1B).

**Figure 1.**
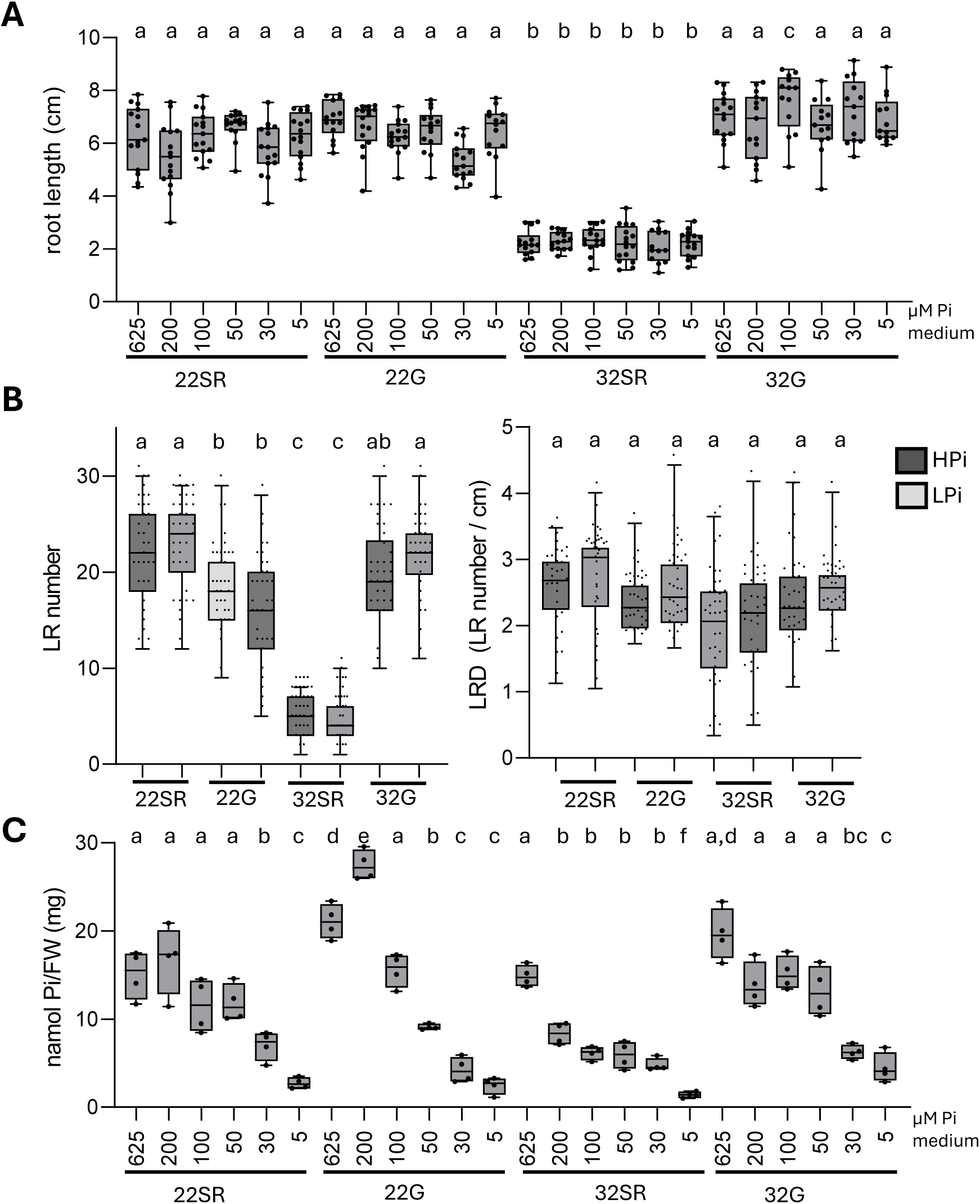
Phosphate (Pi) level in the medium affects root growth and phosphate accumulation in Arabidopsis. (**A**) Primary root length of Arabidopsis seedlings grown at 22°C homogeneous root and shoot (22SR), 22°C shoot and a gradient temperature in the root (22G), 32°C homogeneous root and shoot (32SR), 32°C shoot and a gradient temperature in the root (32G) for 6 days after transfer to media supplemented with 625, 200, 100, 50, 30 or 5 μM of inorganic phosphate (Pi). Boxes represent the interquartile range, the centre line the median and whiskers extend to the most extreme values within 1.5 × IQR (Tukey); dots show individual seedlings (n ≥ 18). **(B)** Number of lateral roots (LRs) and lateral root density (LRD) of seedlings grown for 4 days at 22°C homogeneous in high Pi (625 μM, HPi) or low Pi (30 μM, LPi) and then transferred for 6 days to the different temperature regimes. **(C)** Shoot phosphate content, expressed as nanomoles Pi per mg fresh weight (FW), measured in the same seedlings as in (A) grown under the indicated Pi concentrations. Different lowercase letters above boxplots indicate statistically significant differences among treatments (ANOVA followed by Tukey’s post hoc test, P < 0.05).

Since low temperature can reduce root growth (Zhu et al., 2014), we measured daily root growth after transfer of four-day-old seedlings to the four temperature regimes under HPi or LPi, to rule out an effect of the cooler part of the gradient. From day 2 post-transfer, daily root growth was significantly reduced in 32SR seedlings, which arrested growth almost completely by days 5–6 (Fig. S2). In contrast, 32G seedlings grew at rates similar to those of 22G seedlings irrespective of Pi availability, and both gradient regimes grew slightly less than 22SR (cumulative growth over 8 days of c. 9–10 cm vs. 10.6–11.8 cm; Fig. S2A). Pi deficiency reduced daily root growth only at 22SR, between days 2 and 5 after transfer (Fig. S2A). Thus, the cooler root zone of the TGRooZ only moderately reduces root growth, whereas homogeneous heating of the whole seedling (32SR) arrests it.

We next analyzed shoot Pi content in seedlings grown at different Pi concentrations (Fig. 1C). Homogeneous 32°C significantly reduced Pi accumulation compared with 32G at most Pi concentrations below 625 µM, whereas 22G seedlings showed higher Pi levels at 625 and 200 µM. Pi levels in 32G seedlings were comparable to those at 22SR or 22G, suggesting that defects in root functionality caused by excessive heat in 32SR are responsible for the altered Pi homeostasis.

### Phosphate starvation and heat stress disrupt plant nutrient homeostasis

We next analyzed the effect of the individual and combined stresses on nutrient homeostasis in roots and shoots. Clustering of elemental profiles indicated that heat exerted a stronger influence than Pi starvation in both organs (Fig. S3A,B). In shoots, Pi starvation significantly decreased the levels of P, Mn and Ca, whereas heat increased Na and decreased S, Mg, Mn, Ca and Si (Fig. 2A). The combined stress intensified these changes for most elements, increasing Na and significantly reducing K, S, Zn, P, Ca, Si, Mg and Mn, whereas Mo, B and Fe shifted towards control values (Fig. 2A). In roots, Pi deficiency significantly reduced K, Fe, Si and Zn, with a slight, non-significant increase in Ca, while heat significantly reduced S, K, Fe, Si, Mg and Zn and slightly increased P (Fig. 2B). The combined stress markedly increased Ca and reduced all other elements except P (significantly S, Na, K, Fe, Si, Mg and Zn). Notably, Mn was below the detection limit in roots under all conditions except the combined stress (Fig. S3B). Thus, each individual stress elicits a specific ionomic profile that differs from that of the combined stress.

**Figure 2.**
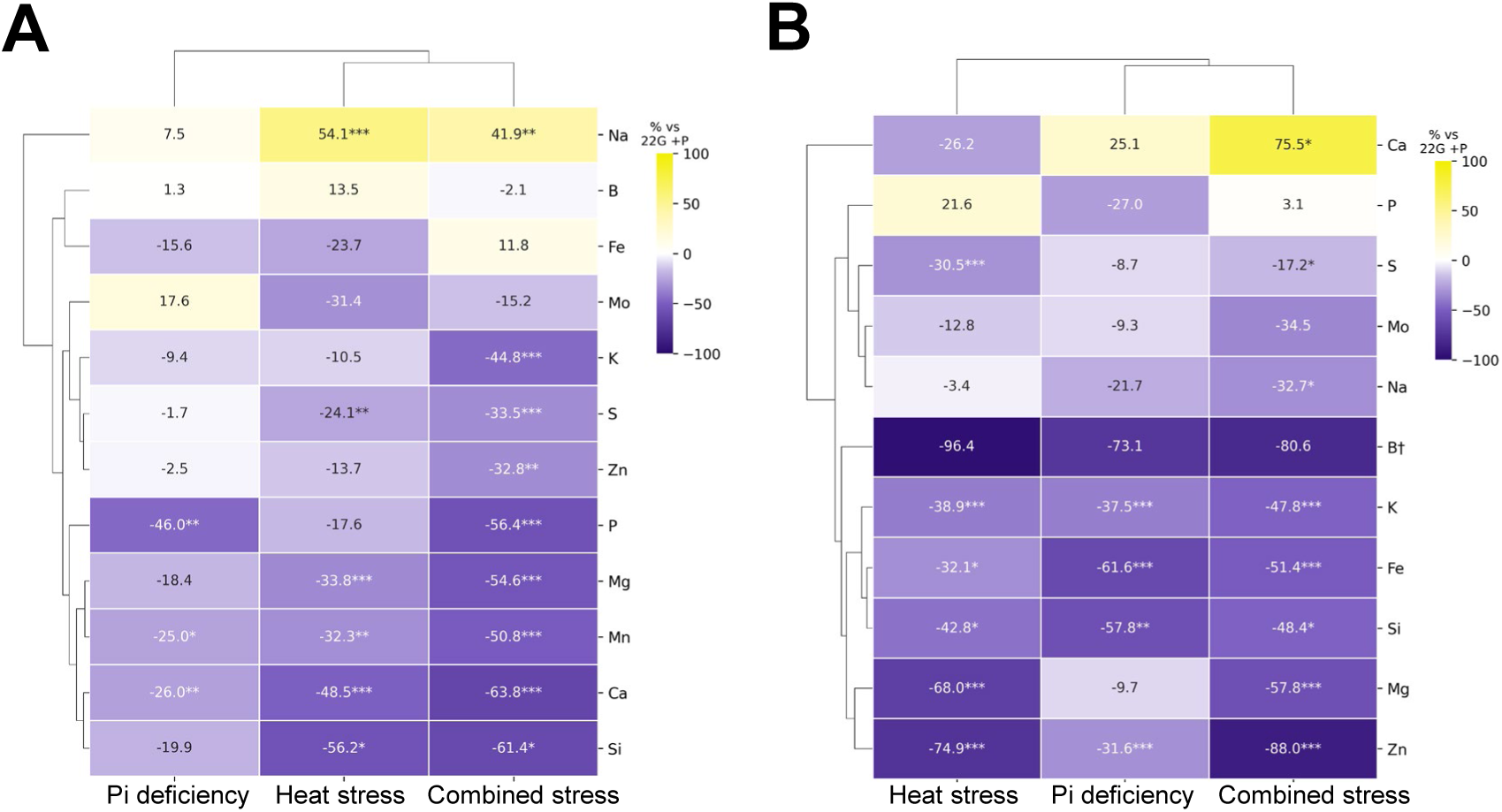
Phosphate starvation and heat stress alter elemental composition in Arabidopsis shoots and roots. Ionomic changes in shoot (A) and root (B) tissue under phosphate (Pi) deficiency, heat stress, and combined Pi deficiency and heat stress. Heatmaps show the percentage change in elemental concentration for each treatment relative to control plants, defined as 22G-HPi. Color scale: yellow indicates an increase and purple a decrease relative to control, ranging from –100% to +100% or higher, exceeding the scale. Elements are hierarchically clustered by their response pattern across treatments. Asterisks denote statistical significance of each treatment versus control (*, p < 0.05; **, p < 0.01; ***, p < 0.001; one-way ANOVA with Dunnett post-hoc correction). Data represent means of n = 4 biological replicates per treatment. †, level below the detection limit in some samples.

### The combination of phosphate starvation and heat triggers a specific gene expression response

We previously analyzed the effect of high temperature on Pi deficiency at the transcriptomic level, focusing on the relationship between DNA methylation and transcription (Lozano-Enguita *et al*., 2026). Here, we focused on genes involved in heat stress, Pi starvation, metabolism or hormone signaling. Clustering of differentially expressed genes (DEGs, Table S2) revealed that Pi starvation had a stronger influence than heat on transcription in roots, whereas the opposite was observed in shoots (Fig. 3A,B). Heat stress and Pi deficiency elicited both common and distinct responses, and the combined stress partially overlapped with the individual stresses but also triggered unique transcriptional changes (Fig. 3C,D). The heatmaps revealed sets of genes whose expression increased (red rectangles) or decreased (green rectangles) under the combined stress relative to both the control and the individual stresses (Fig. 3A,B). GO analysis of the induced clusters (1a and 2a in roots; 1b and 2b in shoots) revealed diverse functional categories (Fig. 3E–H), among which ‘response to jasmonic acid’ and/or ‘response to wounding’ (largely mediated by JA) were represented in all clusters. JA has been linked to Pi starvation (Khan et al., 2016; Li et al., 2024; Sun et al., 2023) and to heat stress (Pandey et al., 2025; Wang et al., 2021; Yang et al., 2019), but its role in the response to their combination remains unknown. Analysis of JA-related genes (Table S3) identified several groups whose expression was enhanced or decreased by the combined stress (Fig. S4A), suggesting that JA signaling might mediate the adaptation of Arabidopsis to combined Pi deficiency and heat.

**Figure 3.**
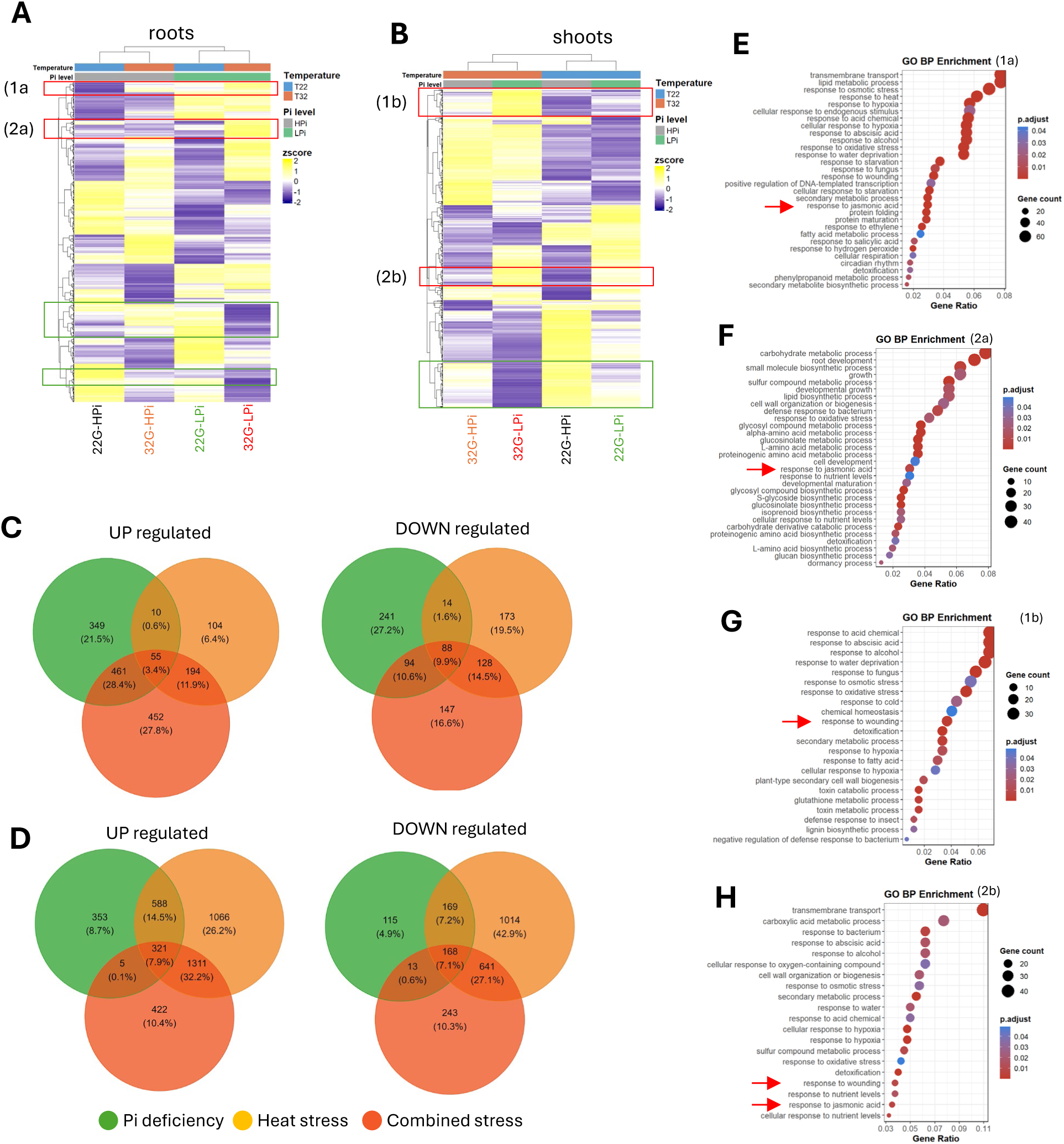
Heat and Pi starvation trigger distinct gene expression patterns relative to combined stress. **(A–B)** Heat maps of genes from seedlings subjected to heat stress (32G-HPi), Pi starvation (22G-LPi), or the combination of both stresses (32G-LPi) in root (A) and shoot (B). The heat maps were clustered by gene expression and treatment. **(C–D)** Venn diagrams showing up- and down-regulated differentially expressed genes (DEGs) in response to heat (32G-HPi vs 22G-HPi), Pi starvation (22G-LPi vs 22G-HPi), or combined stress (32G-LPi vs 22G-HPi) in roots (C) and shoots (D). **(E–F)** Gene ontology of the DEGs in clusters 1a (E) and 2a (F) selected in A. **(G–H)** Gene ontology of the DEGs in clusters 1b (G) and 2b (H) selected in B. Numbers indicate clusters with distinct regulation under combined stress relative to the individual stresses. Red arrows indicate GO terms related to jasmonate.

We also examined genes involved in heat stress responses or Pi homeostasis (Table S3). In both organs, a set of heat-induced genes showed even higher or lower expression under the combined treatment (Fig. S4B). In roots, most Pi starvation-induced genes tended to be less expressed at high temperature, consistent with the slight increase in root P under heat (Fig. 2B), although a specific subset was increased or decreased under combined stress relative to the control and to both individual stresses (Fig. S4C).

To identify responses specific to the combined stress, we extracted genes with a significant Heat × Pi interaction in a two-factor model, i.e. genes whose response cannot be explained by the additive effects of each stress alone, and compared their enriched GO terms with those of heat- and Pi deficiency-responsive DEGs (Table S4; Fig. S5). In roots, the interaction enhanced ion homeostasis and response to iron and repressed processes related to heat and cellular response to Pi starvation. In shoots, it triggered a largely specific program, inducing catabolism, water status and nitrogen recycling and repressing defense, immune responses and mitochondrial signaling.

### Phosphate starvation combined with heat affects plant hormone levels

Given the enrichment of JA-related terms, we analyzed hormone levels under the individual and combined stresses. In shoots, JA decreased under all stresses, although significantly only under the combined stress, while JA-Ile only tended to increase under Pi deficiency and heat (Fig. 4A). In roots, JA increased in all treatments, although significantly only under the combined stress, while JA-Ile increased significantly under all stresses, reaching the highest levels under the combined stress (Fig. 4B). Clustering of hormone profiles indicated that Pi deficiency had a stronger influence than heat in both organs (Fig. 4; Fig. S3C,D). In shoots, Pi deficiency significantly increased cZR and iPR and reduced tZ and its conjugated form tZR, whereas heat significantly increased SA, decreased ABA, iPR and tZR, and slightly increased IAA (Fig. 4A). The combined stress reduced all hormones analyzed, most strongly iPR, SA, tZ, JA and tZR (Fig. 4A). In roots, Pi starvation significantly increased JA-Ile, cZR, iPR, ABA and cZ and reduced tZ, tZR, IBA and IAA, while heat increased JA-Ile, JA and iP and strongly reduced cZR, iPR, tZR, cZ, ABA and IBA (Fig. 4B). The combined stress significantly increased JA-Ile and JA and reduced cZR, iPR, tZR, IBA, IAA, ABA and tZ (Fig. 4B). As previously reported (Silva-Navas *et al*., 2019), Pi starvation increased cZ and reduced tZ in roots, whereas heat reduced cZ without affecting tZ. Under combined stress, cZ remained almost unaffected while tZ decreased to the level of Pi-starved seedlings, suggesting that the stress combination suppresses the Pi deficiency-specific cZ/tZ response. Thus, each individual stress elicits a specific hormonal profile that differs from that of the combined stress.

**Figure 4.**
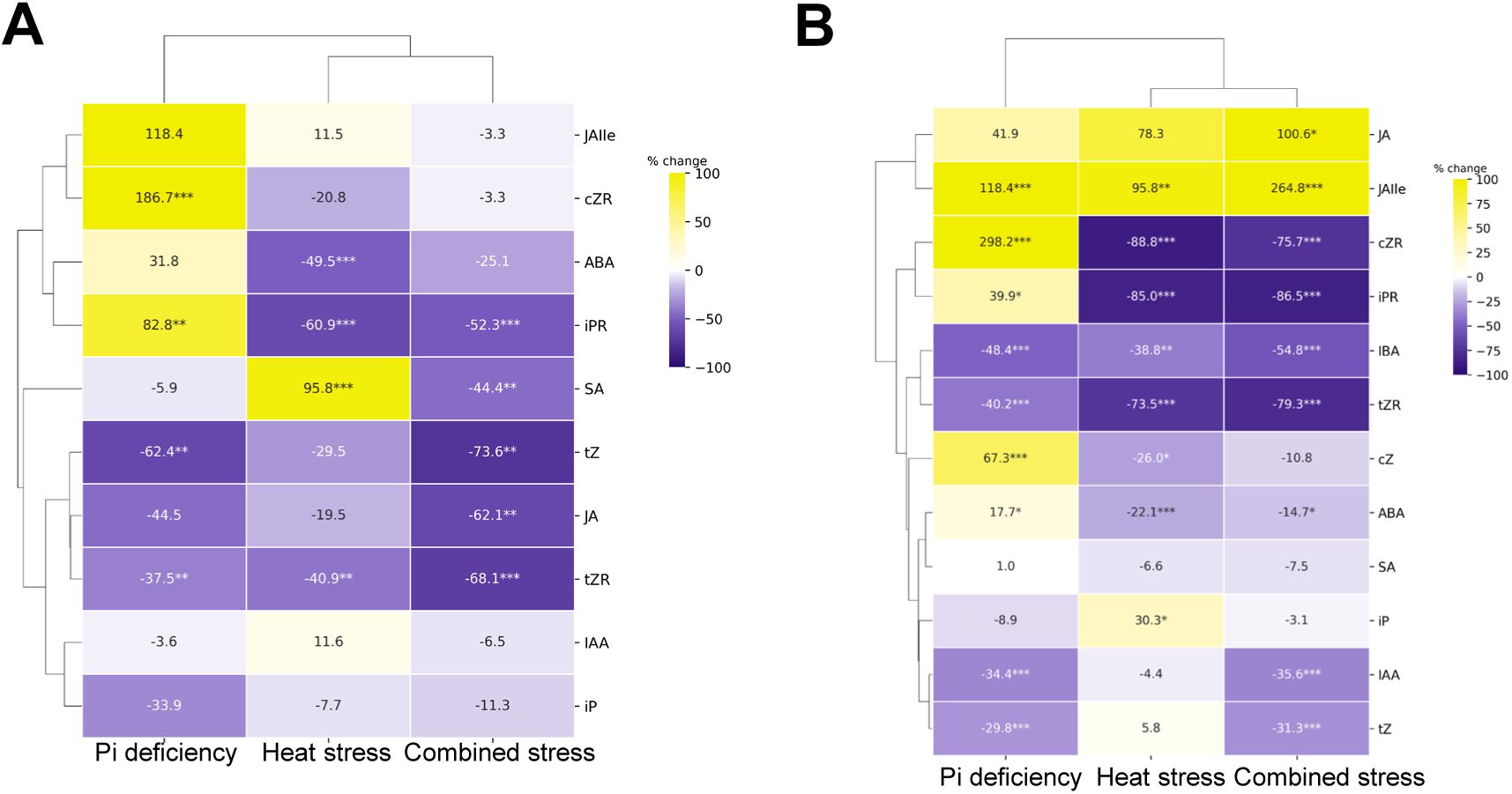
Phosphate starvation and heat stress alter hormone levels in shoots and roots. Heatmaps show the percentage change in hormone content for each treatment, phosphate deficiency (Pi deficiency), heat stress, and combined stress (Pi deficiency plus heat stress) relative to control plants (22G-HPi) in shoots **(A)** or roots **(B)**. Color scale: yellow indicates an increase and purple a decrease relative to control, ranging from –100% to +100% (or higher, where values exceed the scale). Hormones are hierarchically clustered by their response pattern across treatments (dendrograms, top and left). Data were log10-transformed and analyzed by one-way ANOVA followed by Dunnett’s test versus control, with p-values adjusted for 3 comparisons. Asterisks denote statistical significance (*, p < 0.05; **, p < 0.01; ***, p < 0.001). Data represent means of n = 4 biological replicates per treatment.

### JA signaling modulates the transcriptional response to combined phosphate starvation and heat stress

As JA-Ile strongly accumulated in roots under combined stress, we analyzed the transcriptional response of WT seedlings and the *jar1-1* mutant, which has reduced levels of the bioactive form JA-Ile (Staswick *et al*., 2002; Suza and Staswick, 2008). Both genotypes displayed a similar number of DEGs in response to the different stresses (Fig 5A-C, Table S5). In both, the combined stress shared DEGs with the individual stresses in roots and shoots, although combined stress-specific DEGs accounted for a high percentage (Fig. 5A,B). Transcriptional responses were partially conserved between WT and *jar1-1* (44.5–67.9% of up-regulated and 31.0–62.2% of down-regulated DEGs shared). The exception was the root response to heat, where only 18.1% of up- and 19.9% of down-regulated DEGs were shared. Under this condition, *jar1-1* displayed nearly twice as many DEGs as WT (665 vs. 366 up- and 334 vs. 196 down-regulated genes), and *jar1-1*-specific genes accounted for 58.1% and 55.7% of up- and down-regulated DEGs, respectively (Fig. 5C), suggesting that JA-Ile signaling may restrain the root transcriptional response to heat. In addition, overlap was generally lower for down-than for up-regulated DEGs, particularly in roots under combined stress (38.6% vs. 61.3%) and in shoots under heat (31.0% vs. 44.5%). These results suggest that JA-Ile mainly modulates the root transcriptional response to heat and shapes gene repression under stress. Hierarchical clustering of genes across control and combined-stress conditions revealed distinct expression patterns between WT and *jar1-1* in roots, whereas in shoots the overall pattern was largely similar between genotypes (Fig. 5D,E). In roots, two gene sets were specifically induced (set 1d) or repressed (set 2d) under combined stress in *jar1-1*, while these changes were largely absent or attenuated in WT plants (Fig. 5D). In shoots, most genes followed a shared trajectory in both genotypes, with smaller sets showing weaker genotype-specific induction (1e) or repression (2e,3e) (Fig. 5E).

**Figure 5.**
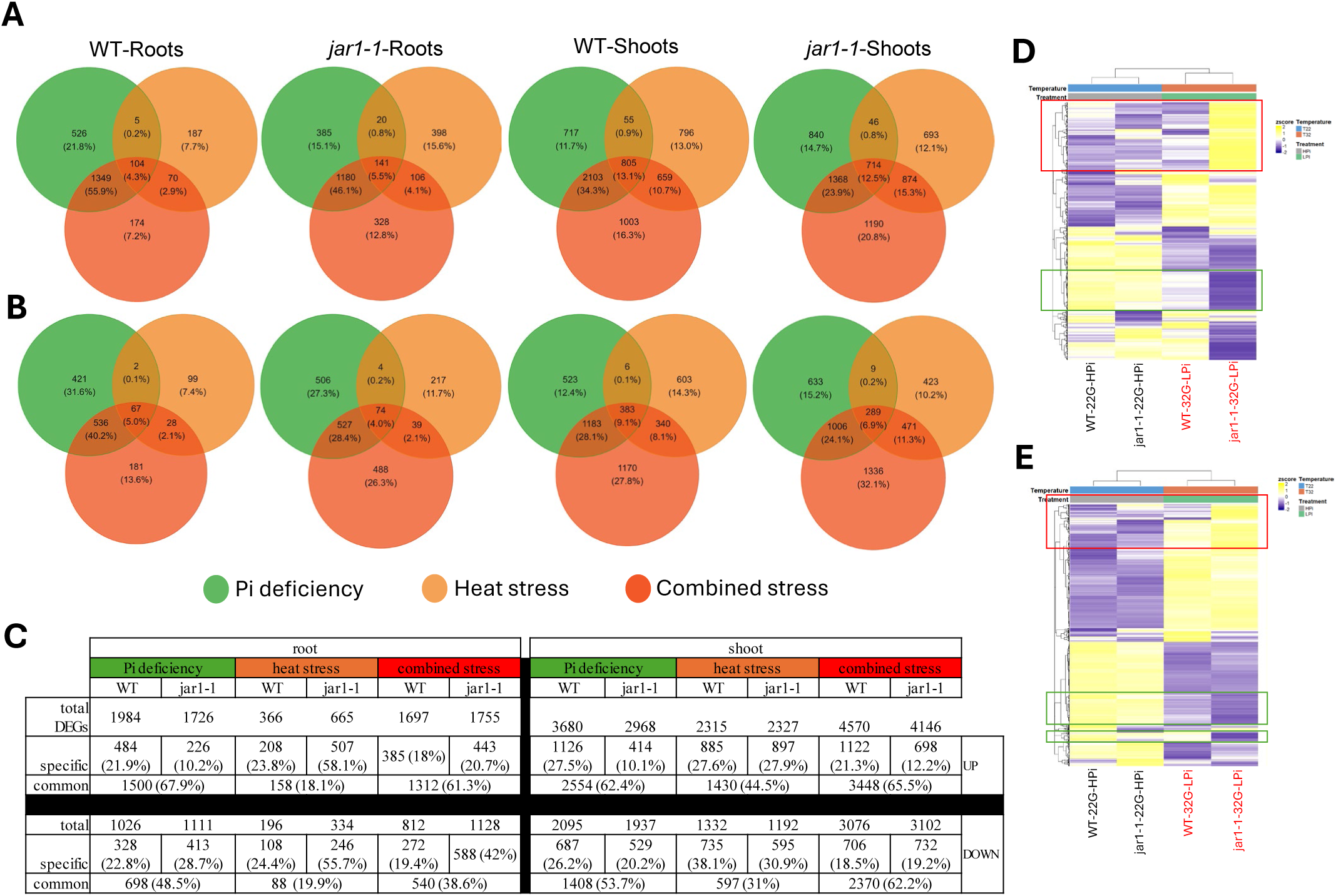
*JAR1*-dependent signaling shapes the transcriptional response to combined heat and Pi starvation. (A–B) Venn diagrams showing the overlap of up-regulated **(A)** and down-regulated **(B)** differentially expressed genes (DEGs) in roots and shoots of wild-type (WT, Col-0) and *jar1-1* seedlings in response to Pi deficiency (22G-LPi vs. 22G-HPi; green), heat stress (32G-HPi vs. 22G-HPi; orange) and combined stress (32G-LPi vs. 22G-HPi; red). DEGs were identified with DESeq2 (adjusted p < 0.05; |log₂ fold change| > 0.75). Numbers indicate the number of genes in each sector, and percentages, in brackets, are relative to the total number of DEGs in each diagram. **(C)** Table showing the specific or common DEGs (absolute value or percentage (%)) in response to Pi deficiency, heat stress or combined stress in roots and shoots between WT and *jar1-1*. **(D–E)** Heatmap of gene expression in WT or *jar1-1* mutant in control condition (22G-HPi) or combined stress (32G-LPi) in roots (D) or shoots (E). Red rectangles indicate sets of genes induced by the combined stress and green rectangles sets of genes repressed.

To better understand the role of JA-Ile, we compared GO terms enriched among Heat × Pi interaction genes in WT and *jar1-1* (Fig. 6; Table S6). The effect of the *jar1-1* mutation was strongly organ-dependent. In roots, the interaction response was smaller than in shoots and slightly reduced in *jar1-1* (110 vs. 159 genes; Fig. 6A,B), possibly because roots grown in the TGRooZ were protected from excessive heat. In both genotypes, the interaction promoted responses to nutrient levels and branched-chain amino acid and glucosinolate biosynthesis, and repressed cellular responses to Pi starvation and oxidative stress. Only WT roots induced Mn and Fe homeostasis, nitrate assimilation and a subset of Pi starvation responses, and repressed responses to temperature, hypoxia and biotic stress, interaction with microorganisms and Pi transport. In shoots, the number of interaction genes was about three times higher in *jar1-1* than in Col-0 (2,458 vs. 830). Both genotypes repressed defense-related processes, including immune response, salicylic acid signaling and programmed cell death (Fig. 6D), but this repression was much broader in *jar1-1*, which showed 94 additional down-regulated terms related to wounding, fatty acid responses, regulation of JA-mediated signaling and callose metabolism. No up-regulated GO terms were shared (Fig. 6C): WT shoots induced catabolic processes, water transport and response to water deprivation, hormone responses, phenylpropanoid biosynthesis and nitrogen recycling, whereas *jar1-1* enhanced translation and ribosome biogenesis, mitochondrial respiration, photorespiration and oxidative stress responses. Thus, JA-Ile signaling restrains the shoot transcriptional response to combined stress, whereas in roots it is required for the full response, particularly for metal homeostasis and the repression of heat- and stress-responsive processes.

**Figure 6.**
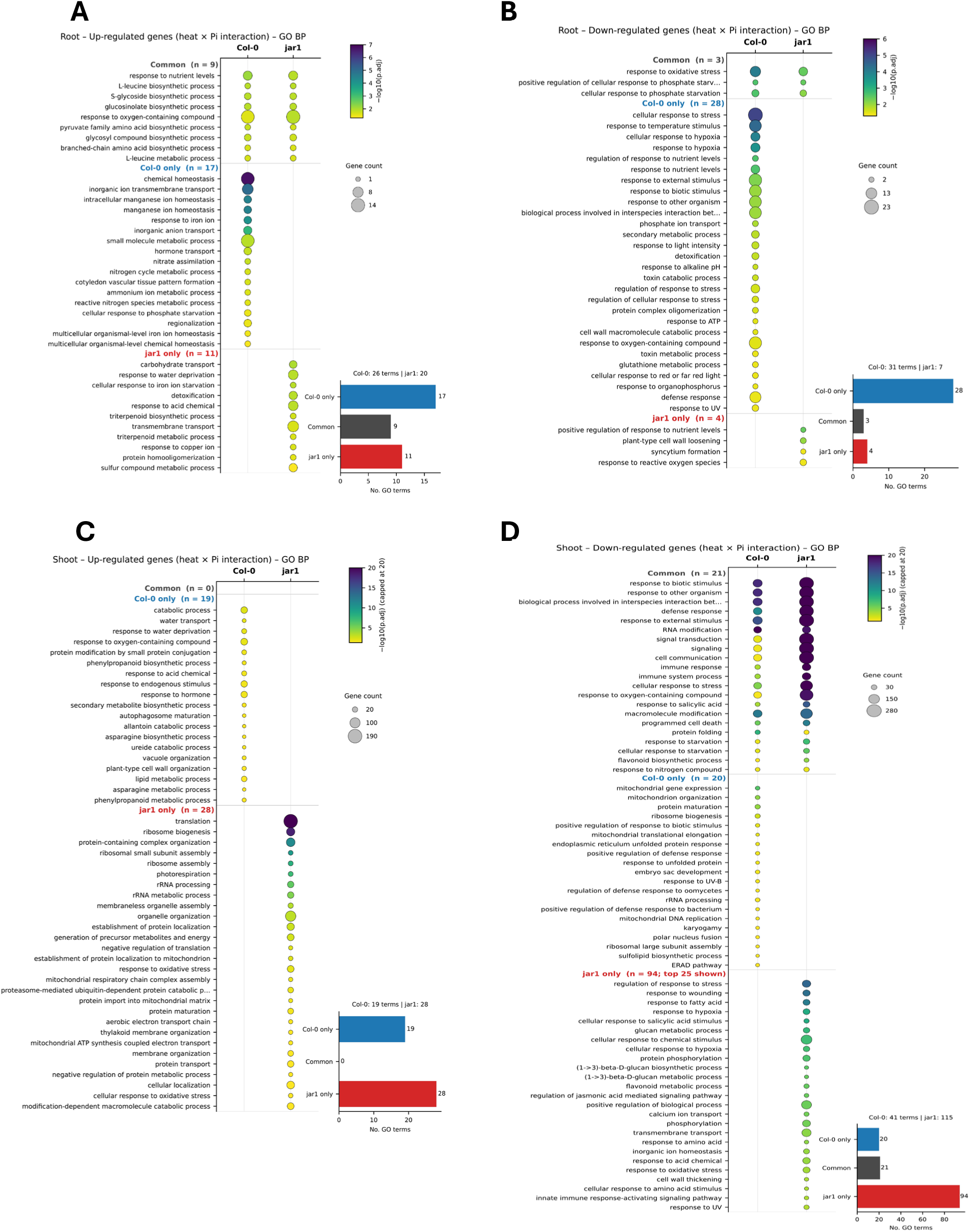
GO biological processes common to or specific for Col-0 and *jar1-1* among genes showing a significant Heat × Pi interaction. GO enrichment of genes with a significant Heat × Pi interaction in wild-type Col-0 and in the JA-Ile-deficient *jar1-1* mutant. Up- and down-regulated interaction genes were analyzed separately. **(A)** Root, up-regulated genes. **(B)** Root, down-regulated genes. **(C)** Shoot, up-regulated genes. **(D)** Shoot, down-regulated genes. Significantly enriched GO terms (adjusted p-value < 0.05) are grouped as common to both genotypes, specific to Col-0 or specific to *jar1-1*, and ordered by significance within each group. In (D), only the 25 most significant *jar1-1-specific* terms are shown (total number indicated in brackets). Dot size is the number of genes annotated to each term, and dot colour is the −log10 of the adjusted p-value; the color scale is panel-specific and, in (C) and (D), capped at 20. Bar plots show the number of GO terms in each category, and the total number of enriched terms per genotype is indicated above them. GO enrichment was performed with clusterProfiler tool using the Arabidopsis genome as background, and p-values were adjusted by the Benjamini–Hochberg method.

The expression of selected heat- and Pi-related genes (Table S3) also differed between WT and *jar1-1* mainly in roots (Fig. S6). Heat-related genes induced under combined stress were generally less expressed in *jar1-1*, whereas a set of genes slightly down-regulated in WT was strongly induced in the mutant; chaperone genes, but not aquaporins, showed a similar pattern. Many Pi-related genes slightly induced in WT were strongly up-regulated in *jar1-1* under combined stress. In addition, a large number of ABA-related genes were differentially regulated in WT and *jar1-1* under combined stress (Fig. S6), suggesting an interconnection of JA and ABA signaling. Together, these results support a role for JA-Ile in coordinating the transcriptional response to combined heat and Pi deficiency.

### JA signaling modulates plant growth under combined heat and Pi starvation in vitro and in soil

To test the physiological role of JA, we analyzed two mutants with opposite JA signaling: *jar1-1*, with lower JA-Ile content, and *jazQ*, a quintuple mutant lacking the *JAZ1, JAZ3, JAZ4, JAZ9,* and *JAZ10* transcriptional repressors, which shows de-repressed JA signaling. We first grew these mutants and WT under HPi or LPi at 22G or 32G in ½ MS medium (Fig. 7A–D). Primary root growth of WT was largely unaffected by Pi deficiency, heat or their combination, whereas *jar1-1,* which had shorter roots than WT under HPi, recovered WT-like root length under LPi at both temperatures (Fig. 7A). In contrast, *jazQ* seedlings, which already had a shorter primary root than WT or *jar1-1*, showed a significant reduction in root growth under LPi, heat and combined stress (Fig. 7A). LR number was significantly reduced in WT and *jazQ* only under combined stress, and in *jar1-1* under heat and combined stress (Fig. 7B). Under HPi, *jar1-1* produced more and *jazQ* less shoot fresh weight than WT, and 32G reduced shoot fresh weight in *jar1-1* and *jazQ* but not in WT (Fig. 7C). Under LPi, heat significantly reduced shoot fresh weight in WT (Fig. 7D). Relative to 22G-HPi, combined stress reduced biomass by 61.9% in WT, 62.5% in *jar1-1* and 47.6% in *jazQ*, which may partly reflect the lower baseline growth of *jazQ*.

**Figure 7.**
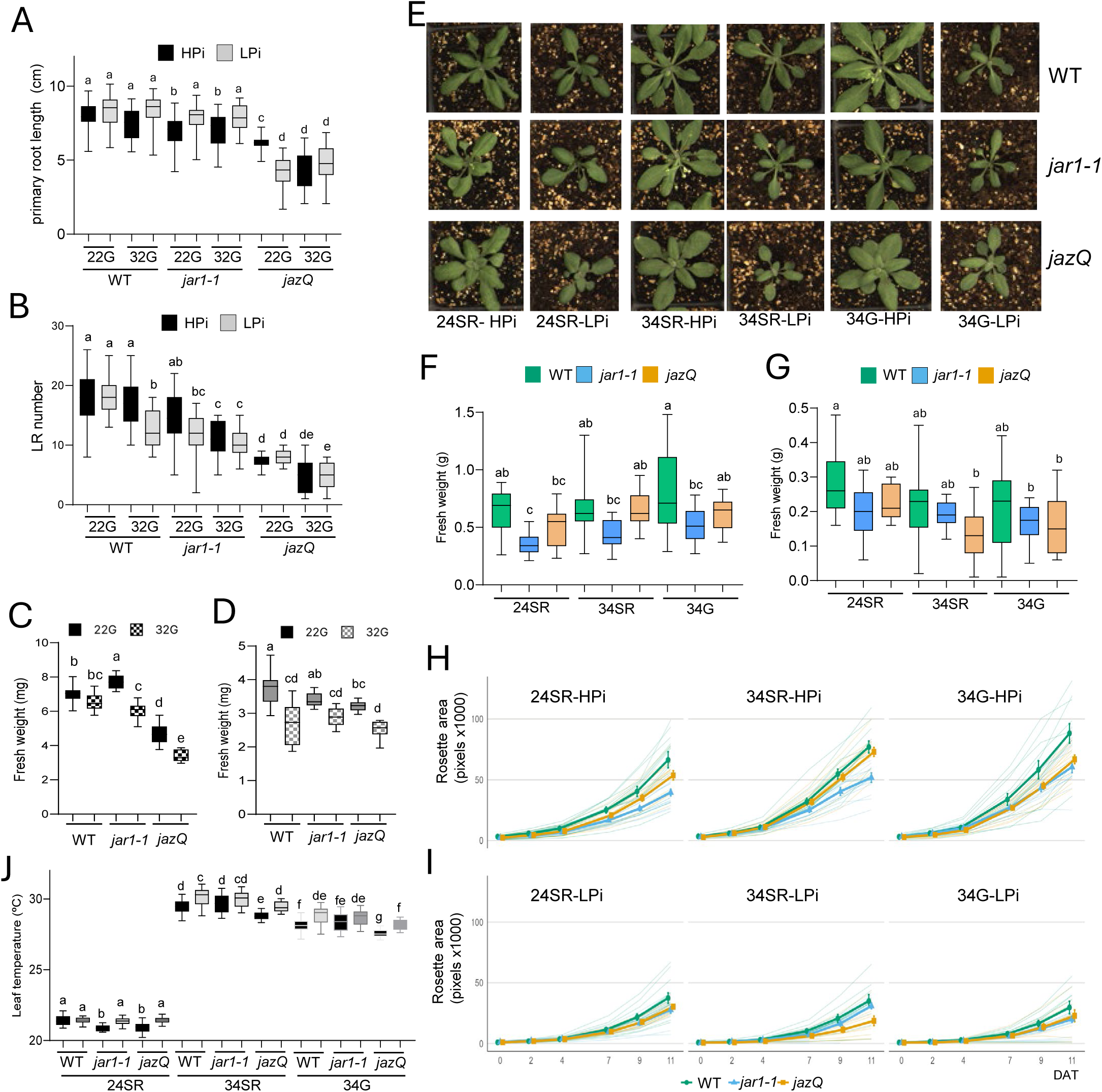
Heat stress and phosphate deficiency elicit shared and distinct phenotypic responses in vitro and in soil. **(A–D)** WT, *jar1-1* and *jazQ* seedlings were grown on ½ MS medium containing high Pi (HPi) or low Pi (LPi) for 4 days at 22 °C in the TGRooZ system (22G). Half of the plates were then kept at 22G and the other half were transferred to 32 °C in the TGRooZ (32G) for 6 days. n ≥ 20. **(A)** Primary root length and **(B)** lateral root (LR) number under HPi (black boxes) or LPi (grey boxes). **(C–D)** Shoot fresh weight under HPi (C) or LPi (D) at 22G (black boxes) or 32G (checkered boxes). **(E–J)** WT, *jar1-1* and *jazQ* seedlings were grown on ½ MS medium with HPi or LPi for 7 days at 22 °C, transplanted into 0.5-L pots containing soil with high or low Pi level, and acclimated at 24 °C for 2 days. Plants were then transferred to 24 °C homogeneous temperature (24SR), 34 °C homogeneous temperature (34SR) or 34 °C in the TGRooZ (34G) for 11 days. Days after transfer (DAT) refer to the start of the temperature treatments. n = 13 per genotype. **(E)** Representative images of plants at 9 DAT. **(F–G)** Rosette fresh weight at 11 DAT under HPi (F) or LPi (G). **(H–I)** Rosette growth dynamics under HPi (H) or LPi (I), quantified from top-view images as projected rosette area (pixels × 1000) at 0, 2, 4, 7, 9 and 11 DAT. Thick lines show mean ± SD; thin lines show individual plants. **(J)** Leaf temperature of plants grown under HPi (black boxes) or LPi (grey boxes) at 24SR, 34SR or 34G, measured *by* infrared thermography camera at 6 DAT. In box plots, boxes represent the interquartile range, the centre line the median and whiskers the minimum and maximum values. Different letters indicate significant differences within each panel (one-way ANOVA followed by Tukey’s HSD test, *P* < 0.05).

We next analyzed these genotypes in soil fertilized with HPi or LPi (see Materials and Methods) at 24 °C (24SR), homogeneous 34 °C (34SR) or 34 °C in the TGRooZ (34G) (Fig. 7E–I). In contrast to in vitro, under control conditions (24SR-HPi) *jar1-1* showed 44% lower rosette fresh weight than WT, whereas *jazQ* did not differ significantly from WT (Fig. 7F). Under HPi, heat did not reduce biomass in any genotype and even tended to increase it, particularly at 34G (+24%, +47% and +23% in WT, *jar1-1* and *jazQ*, respectively; Fig. 7F). Relative to their 24SR-HPi controls, Pi deficiency reduced rosette fresh weight by 44–55%, and heat aggravated this reduction to ∼65% in WT, 46–54% in *jar1-1* and 67–73% in *jazQ* (Fig. 7F,G). Accordingly, a three-way ANOVA revealed a significant temperature × Pi interaction (P < 0.001), indicating that heat compromises shoot growth mainly when Pi is limiting.

Rosette growth dynamics were consistent with the fresh weight data (Fig. 7H,I). Under HPi, rosette area increased exponentially from 4 DAT in all genotypes, and genotypic differences became evident from 7 DAT onwards. In contrast to in vitro biomass, *jar1-1* displayed the slowest rosette growth under all temperature regimes, reaching ∼40% less area than WT at 11 DAT at 24SR, whereas *jazQ* showed intermediate values (Fig. 7H). Heat did not impair rosette expansion under HPi; rather, WT rosette area increased by ∼15% and ∼30% at 34SR and 34G, respectively. Pi deficiency strongly restricted rosette growth in all genotypes (∼40% reduction in WT at 24SR) and attenuated genotypic differences (Fig. 7I). Under LPi, heat no longer promoted growth and instead tended to reduce WT rosette area, particularly at 34G, whereas *jazQ* was most affected at 34SR, reaching about half of the WT rosette area at 11 DAT (Fig. 7I).

Finally, we measured leaf temperature at 6 DAT (Fig. 7J). At 24SR, WT leaf temperature did not respond to LPi, whereas *jar1-1* and *jazQ*, which had cooler leaves than WT under HPi, increased it to WT levels under LPi. Heat increased leaf temperature in all genotypes, although leaves were ∼1 °C cooler at 34G than at 34SR, and *jazQ* showed the lowest leaf temperature under HPi in both heat regimes. Under heat, LPi further increased leaf temperature in WT and *jazQ*, but not significantly in *jar1-1*. These results suggest that JA signaling and Pi status interact in the regulation of growth and leaf temperature under heat.

## Discussion

Plants in natural environments rarely experience stresses in isolation. Elevated temperature combined with nutrient limitation represents a major challenge for crop productivity and food security (Hatfield and Prueger, 2015; Köhler et al., 2019), and its negative effects are expected to be exacerbated by climate change. Using the TGRooZ system to mimic natural soil conditions (González-García *et al*., 2022), we show that combined Pi starvation and heat elicit distinctive nutritional, hormonal and transcriptional responses that differ from those to each individual stress, and identify JA signaling as a modulator of plant adaptation to this stress combination.

### Root temperature is a major determinant of phosphate homeostasis under heat stress

High temperatures alter root development and nutrient acquisition, often changing plant mineral composition (Köhler *et al*., 2019). As previously observed (González-García *et al*., 2022), homogeneous exposure of shoot and roots to 32°C strongly inhibited root growth and reduced shoot Pi accumulation. However, when roots were protected from excessive heating in the TGRooZ, plants maintained root growth and Pi levels comparable to control conditions. Thus, the reduction in shoot Pi under homogeneous high temperature largely results from impaired root function rather than from altered shoot metabolism. This is particularly relevant under field conditions, where soil temperatures often remain lower than air temperatures and may buffer the negative effects of heatwaves on nutrient homeostasis.

Limitation of one macronutrient alters the plant’s overall nutrient balance, affecting the uptake, translocation and accumulation of other elements (Ågren et al., 2012; Ding et al., 2017; Fan et al., 2021). Similarly, heat stress affects the uptake, transport and assimilation of multiple nutrients (Giri *et al*., 2017; Mishra *et al*., 2023) by modifying membrane fluidity, transporter activity, transpiration and shoot architecture (Zhao *et al*., 2020). In our study, heat exerted a stronger influence on elemental composition than Pi deficiency, and each stress imposed a distinct nutritional signature, with stronger imbalances in shoots than in roots. Pi deficiency caused relatively targeted changes, reducing P, Mn and Ca in shoots and Fe, Zn, K and Si in roots, consistent with the crosstalk between Pi and Fe/Zn homeostasis (Fan *et al*., 2021). Heat mainly depleted divalent cations (Mg, Mn and Ca in shoots; Mg and Zn in roots) and Si, while increasing Na in shoots, which may reflect heat-induced alterations in transpiration and root uptake. The combination of both stresses caused a broad and largely exacerbated depletion of macro- and micronutrients, which might be relevant for designing fertilization programs for soils with low P content and predictable heat waves. Mn was strongly depleted in shoots under combined stress (−51%), whereas in roots it was only detected under this condition, coinciding with the induction of genes related to Mn and Fe homeostasis among Col-0 root interaction genes; this suggests a compensatory transcriptional response and/or impaired root-to-shoot Mn translocation. The reduced Zn, Fe and Mn content under combined stress could be explained by repression of the root Fe-acquisition machinery: *FIT* was down-regulated together with its targets *IRT1*, *AHA2* and the coumarin biosynthetic genes *S8H* and *CYP82C4* (Colangelo and Guerinot, 2004; Rajniak et al., 2018). Because IRT1 also transports Mn²⁺ and Zn²⁺ (Vert et al., 2002), its repression offers a single mechanism for the decline of all three micronutrients. In addition, the strong down-regulation of *NAS1* and *NAS2* suggests reduced nicotianamine biosynthesis, which limits Fe and Zn translocation (Klatte et al., 2009). Meanwhile, internal deficiency markers such as *ZIP1*, *ZIP4*, *IRT3*, *bHLH100* and *IMA2/3* were induced (Assunção et al., 2010; Grillet et al., 2018), suggesting that Pi-status signals override the Fe/Zn-deficiency response at the level of FIT/IRT1, uncoupling deficiency signaling from uptake.

Ca was the only element that accumulated under combined stress, and only in roots. Calcium is a central second messenger in heat sensing, mediating Ca²⁺ influx through CNGC channels and the activation of HSF/HSP-dependent thermotolerance (Kang et al., 2023), whereas Pi starvation dampens stress-induced cytosolic Ca²⁺ signals in roots (Matthus et al., 2019). The opposite changes of Ca under combined stress, higher in roots (+75%) but lower in shoot (−63.8%), may reflect an organ-dependent compensatory mechanism to sustain Ca-dependent signaling during heat responses when Pi is limited, although this hypothesis requires further investigation.

Pi deficiency triggers metabolic reprogramming to optimize internal Pi use and root architectural changes that enhance soil Pi foraging (Madison et al., 2023), whereas heat stress activates protective programs to limit cellular damage. When both stresses coincide, these programs likely compete for resources and impose conflicting demands between organs: the shoot bears the heat load and requires Pi to sustain metabolism, whereas the root must acquire and deliver Pi to the shoot. This divergence is reflected in the exacerbated nutrient depletion and the organ-specific Heat × Pi transcriptional responses under combined stress. Heat and Pi deficiency therefore interact at the levels of nutrient uptake, transport and redistribution, and the elements most sensitive to their combination, particularly K, Mg and Zn, provide targets for nutrient-management strategies in nutrient-poor soils exposed to heat.

### The combined stress, heat and Pi deficiency, elicits a unique hormonal signature

Pi starvation modifies phytohormone levels and signaling, shaping physiological, morphological and molecular responses (Lu et al., 2026; Madison *et al*., 2023; Silva-Navas *et al*., 2019). In our conditions, Pi starvation rather than heat was the major driver of hormone changes, particularly in roots. However, the combined treatment generated a hormonal profile that differed substantially from those induced by either stress, supporting that plants integrate these signals rather than simply summing independent responses. Root morphology is shaped by the balance between cytokinin (CK) and auxin (Rivas et al., 2022; Silva-Navas *et al*., 2019; Tiwari et al., 2023), and CK is considered the main systemic hormone regulating shoot/root responses under Pi starvation (Franco-Zorrilla et al., 2005; Lin et al., 2014). We previously showed that an increase in the cis-Zeatin (cZ) / trans-Zeatin (tZ) ratio helps balance root-shoot growth, favoring root growth and adaptation (Silva-Navas *et al*., 2019). Here, high temperature counteracted the cZ/tZ increase, possibly prioritizing shoot responses to potentially lethal heat over the response to Pi starvation, a hypothesis that remains to be tested.

ABA is increasingly recognized as a central integrator of environmental stress responses and may help coordinate nutrient and temperature signaling (Cutler et al., 2010; González-García et al., 2026; Singh and Roychoudhury, 2023; Vishwakarma et al., 2017). We found that ABA levels increased under Pi starvation (significantly only in roots), but were reduced by heat stress and, in roots, by the combined stress. This contrasts with the heat-induced ABA accumulation commonly reported and may be explained by the use of the TGRooZ, which maintains root functionality during heat stress (Devesa-Aranguren et al., 2026; Fonseca de Lima et al., 2021; González-García *et al*., 2022). Analyses in soil-grown plants will be required to fully understand ABA dynamics in response to heat and Pi starvation.

The most striking hormonal changes involved jasmonates. Beyond their roles in defense against herbivores and pathogens, JA and JA-Ile integrate environmental signals with growth regulation (Wang et al., 2026) and participate in responses to nutrient deficiencies, including phosphate starvation, and to heat stress. Several lines of evidence support a role for JA signaling in the response to phosphate starvation under elevated temperature. First, JA-Ile increased significantly in roots under all stress conditions, reaching the highest levels under combined stress, whereas shoot JA decreased significantly under combined stress. Second, genes associated with JA signaling, wound responses and jasmonate-mediated pathways were among the categories specifically enriched under combined stress. Third, JA mutants showed that JA levels and signaling modulate the balance between stress responses and growth.

### Jasmonic acid modulates the combined stress response

Phosphorus supply modulates heat tolerance in several crops. In common bean, increasing P supply improved photosynthetic performance, dry matter partitioning and seed yield under high temperature in acidic soils (Suárez et al., 2025), and in soybean, a high post-flowering P status enhanced tolerance to terminal heat stress (Laira et al., 2023). However, the mechanisms by which Pi availability shapes heat tolerance remain largely unexplored. Our phenotypic analyses reveal that Pi availability is a major determinant of heat tolerance in Arabidopsis. Under Pi-sufficient conditions, WT plants maintained shoot growth at elevated temperature both in vitro and in soil, where heat even tended to promote rosette growth. Under LPi, however, heat reduced WT shoot fresh weight in vitro and aggravated the growth restriction imposed by Pi deficiency in soil, as supported by the significant temperature × Pi interaction. Thus, Pi deficiency compromises the capacity of plants to sustain growth at high temperature, possibly by limiting energy- and phosphate-demanding processes required for thermal acclimation. Non-additive interactions with species-specific outcomes have been reported in other species: combining low P with root-zone warming, a setup comparable to our TGRooZ, reduced maize dry mass by 55.8% (Xia et al., 2021), whereas in soybean P deficiency mainly exacerbated low-temperature stress (Singh et al., 2018), and in sheepgrass heat and Pi deficiency interacted in a complex manner on photochemical performance (Li *et al*., 2018). These differences likely reflect species-specific thermal optima and P-use strategies.

JA has been independently linked with Pi starvation (Khan *et al*., 2016; Li *et al*., 2024; Sun *et al*., 2023) and with heat stress (He et al., 2023; Pandey *et al*., 2025; Wang *et al*., 2021; Yang *et al*., 2019), contributing to transcriptional reprogramming, root architectural responses, heat acclimation and resource allocation, but its role in coordinating responses to both stresses simultaneously was unknown. While many classical phosphate starvation genes showed reduced expression at elevated temperatures, jasmonate-responsive genes remained strongly induced or were specifically activated under combined stress. This suggests that JA participates in a specific adaptive program rather than in a simple overlay of individual stress responses, as described for other stress combinations (Mittler, 2006; Prasch and Sonnewald, 2013; Rasmussen et al., 2013; Zandalinas et al., 2024).

The comparison between *jar1-1* and *jazQ* indicates that the contribution of JA to these responses is not proportional to JA activity. In vitro, both reduced JA-Ile biosynthesis (*jar1-1*) and constitutive de-repression of JA signaling (*jazQ*) abolished the heat tolerance displayed by WT under HPi (−22% and −27%, respectively, vs. −7% in WT), suggesting that an intact and tunable JA pathway, rather than high or low JA output per se, is required to sustain growth at elevated temperature. Under LPi, differences among genotypes were largely attenuated, indicating that Pi limitation overrides the JA-dependent component of heat tolerance.

Relative growth losses must be interpreted in light of each genotype’s baseline. *jazQ* displayed reduced shoot growth, shorter primary roots and fewer lateral roots in vitro, consistent with the growth–defense trade-off associated with constitutive JA signaling (Campos *et al*., 2016; Zhu et al., 2026). Its smaller relative loss under combined stress in vitro (−48% vs. −62% in WT) therefore largely reflects a compressed growth range. Likewise, in soil, the smaller relative loss of *jar1-1* under combined stress (46–54% vs. ∼65% in WT) arises from a 44% lower baseline, and in absolute terms *jar1-1* did not outperform WT, whereas *jazQ* was the most affected genotype (67–73%). *jar1-1* accumulated more biomass than WT in vitro but less in soil, suggesting that the requirement for JA-Ile depends on the growth environment, for instance to cope with biotic or edaphic factors present in non-sterile soil. Consistently, increased JA-Ile restricts growth but enhances drought tolerance, whereas reduced JA-Ile increases drought sensitivity in Arabidopsis (Mahmud et al., 2022).

These phenotypes can be reconciled by an organ-specific model of jasmonate action. Under combined stress, JA and JA-Ile strongly accumulated in roots (+101% and +265%, respectively), whereas JA decreased in shoots (−62%) and JA-Ile remained unchanged (Fig. 4). Thus, the stress combination rewires jasmonate status in opposite directions in the organ that senses Pi availability and in the organ directly exposed to heat. Root JA-Ile accumulation may contribute to the transcriptional reprogramming of the root, consistent with the altered root heat response and the loss of metal homeostasis-related terms in *jar1-1*, whereas the decrease in shoot JA would relieve JA-mediated growth repression and favor thermal acclimation. Within this framework, *jar1-1* would be impaired in the root component of the response, whereas *jazQ*, unable to attenuate JA signaling in the shoot, would have a permanently lowered growth ceiling, explaining why disrupting the pathway in either direction compromises heat tolerance. Hence, JA appears to tune the balance between stress acclimation and growth in an organ- and dose-dependent manner. Measuring jasmonate levels in roots and shoots of these mutants, and manipulating JA specifically in the root zone, will be required to test this model.

Leaf temperature provides additional clues to the interaction between JA and Pi status. At 24SR, *jar1-1* and *jazQ* displayed lower leaf temperatures than WT under HPi, and Pi deficiency raised them to WT levels, whereas WT leaf temperature was insensitive to Pi supply. Under heat, *jazQ* maintained the lowest leaf temperature under HPi, and LPi increased leaf temperature in WT and *jazQ* but not in *jar1-1*. Since leaf temperature largely reflects transpirational cooling, these results suggest that Pi deficiency restricts transpiration in a JA-modulated manner. JA may influence stomatal behavior through ABA-dependent pathways; poplar plants overexpressing JAZ2 show enhanced ABA-induced stomatal closure and increased leaf temperature (Rao et al., 2023), and loss of *JAZ* repressors in *jazQ* would conversely be expected to attenuate stomatal closure, consistent with our observations. However, the lower leaf temperature of *jar1-1* at 24SR and its lack of response to LPi under heat indicate that JA-Ile also contributes to Pi-dependent regulation of transpiration, again pointing to a non-linear relationship. Leaves were also ∼1 °C cooler at 34G than at 34SR, suggesting that the root-zone temperature gradient improves leaf thermoregulation, likely by sustaining water uptake. Importantly, lower leaf temperature did not translate into better growth, since *jazQ* was the most affected genotype under combined stress in soil. Measurements of stomatal conductance and transpiration will be needed to determine whether JA directly affects thermoregulation or whether these differences are secondary to altered growth and Pi status.

## Conclusions

Phosphate starvation and heat stress interact to generate a unique physiological state characterized by extensive nutrient reprogramming, profound hormonal remodeling and a specific transcriptional response. Although root temperature strongly influences phosphate homeostasis under heat stress, the integration of phosphate and temperature signals results in responses that cannot be predicted from either stress alone and should therefore be investigated as a distinct combined-stress condition. Among the regulatory pathways identified, jasmonic acid signaling emerges as an important regulator of adaptation to the combined stress. The enrichment of JA-responsive genes, together with the contrasting phenotypes of *jar1-1* and *jazQ* mutants, supports a model in which organ-specific changes in jasmonate status, increasing in roots and decreasing in shoots, link nutrient status, thermal stress and growth. These findings identify JA signaling as a promising target for improving plant resilience to abiotic stresses under future climate scenarios.

## ACKNOWLEDGMENTS

We thank Roberto Solano and Andrea Chini for providing the *jar1-1* and *jazQ* mutants. We also thank the CBGP’s greenhouse facility for the help provided.

## Data availability

The RNA-seq datasets for the WT response to heat, Pi deficiency and combined stress have been deposited in the NCBI Gene Expression Omnibus (GEO) under accession number GSE327853.

## Supporting Information

**Figure S1.** Determination of the Pi concentration that activates the phosphate starvation response under the different temperature regimes.

**Figure S2.** Daily primary root growth of *Arabidopsis thaliana* seedlings under different temperature regimes and phosphate availability.

**Figure S3.** Phosphate starvation and heat stress alter ion and hormone accumulation in roots and shoots.

**Figure S4.** Differential expression of genes associated with heat stress, phosphate signaling, and jasmonic acid pathways.

**Figure S5.** Gene Ontology (GO) biological processes enriched among genes responsive to heat, Pi deficiency and their interaction in *Arabidopsis thaliana*.

**Figure S6.** Expression of heat-, chaperone-, aquaporin-, Pi- and ABA-related genes in WT and *jar1-1* plants under control and combined stress conditions.

**Table S1**. Substrate nutrient analysis

**Table S2**. Differentially expressed genes in roots and shoots in response to Pi deficiency, heat and combined stress.

**Table S3**. Genes of the selected Gene Ontology categories (JA-, heat-, Pi- and ABA-related genes, chaperones and aquaporins) shown in Figs. S4 and S6.

**Table S4**. Genes regulated by temperature x Pi deficiency interaction in WT plants

**Table S5.** Differentially expressed genes in roots and shoots in response to Pi deficiency, heat and combined stress in WT and *jar1-1*.

**Table S6.** GO terms enriched among Heat × Pi interaction genes in roots and shoots of WT and *jar1-1*.

## Author contributions

J.C.P. and E.C. planned and designed the research. V.B.-G., F.B., A.L.-E., M.L., I.T.G.-M., A.S., S.N.-N., J.M.G.-M., A.M.Z. and M.G.-O. performed the experiments. J.M.G.-M., A.M.Z. and M.G.-O. performed the ionomic and hormonomic analyses. F.B. carried out the automatic phenotyping and analyses. A.L.-E., F.B. and V.B.-G. carried out the bioinformatic analyses of sequencing datasets. J.C.P. and E.C. analyzed the data and wrote and edited the manuscript.

## Declaration of generative AI and AI-assisted technologies in the writing process

During the preparation of this manuscript, the authors used Claude (Anthropic) to improve clarity, structure and language. After using this tool, the authors reviewed and edited the content as needed and take full responsibility for the content of the publication. No AI tools were used for data analysis, interpretation or generation of results.

