## Supplemental Figures for "Integrated multi-omics reveals organ-specific jasmonate signaling in Arabidopsis under combined heat and phosphate starvation"

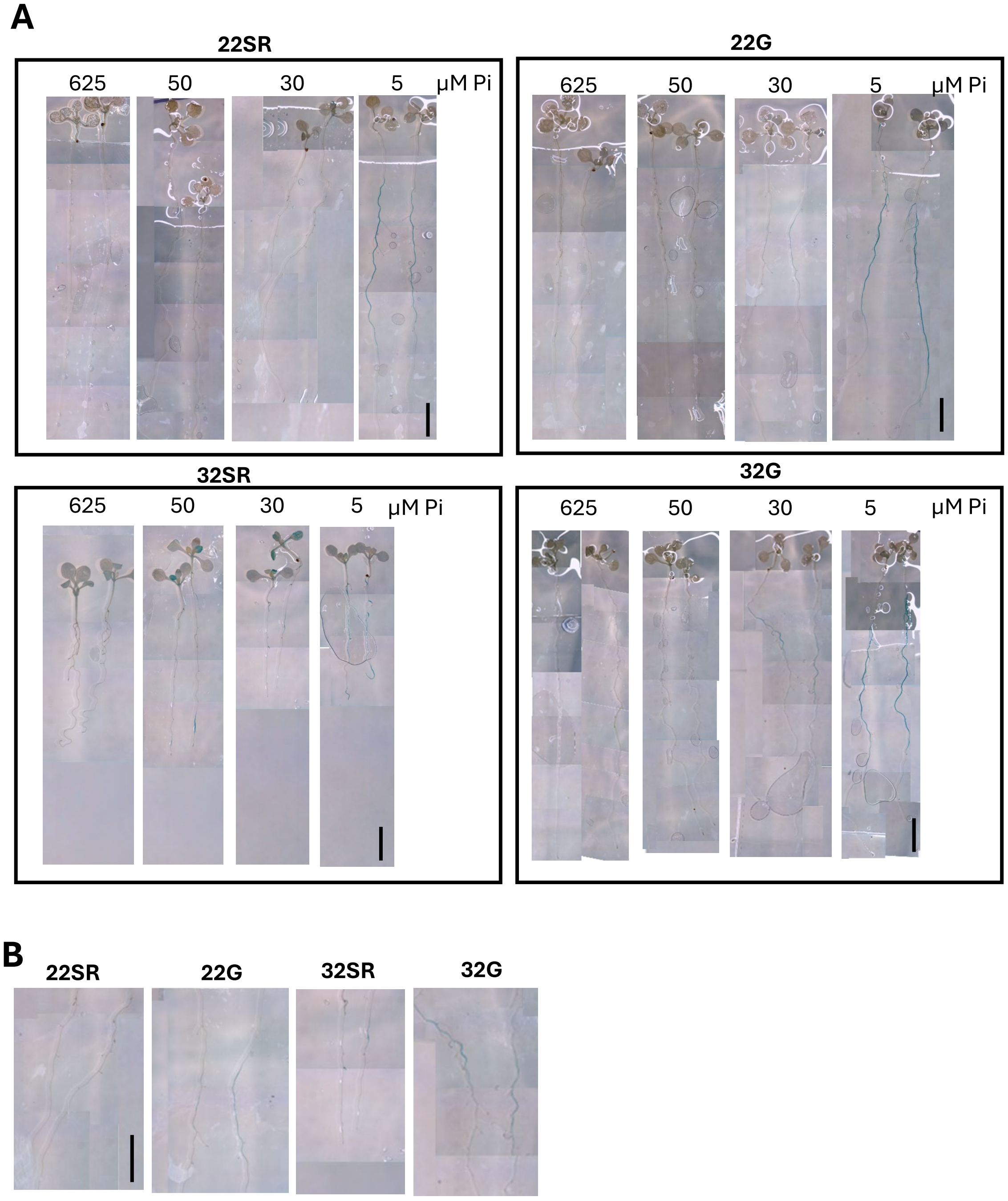


**Figure S1. Determination of the Pi concentration that activates the phosphate starvation response under the different temperature regimes. (A)** GUS staining of *SPX3::GUS* marker line grown at 22ºC homogeneous root and shoot (22SR), 22ºC shoot and a gradient temperature in the root (22G), 32ºC homogeneous root and shoot (32SR), 32ºC shoot and a gradient temperature in the root (32G) for 6 days after transference on media supplemented with 625, 50, or 30 or 5 μM of inorganic phosphate (Pi). **(B)** Higer magnification of the central root section of plants grown and in A in 30 μM of inorganic phosphate (Pi).

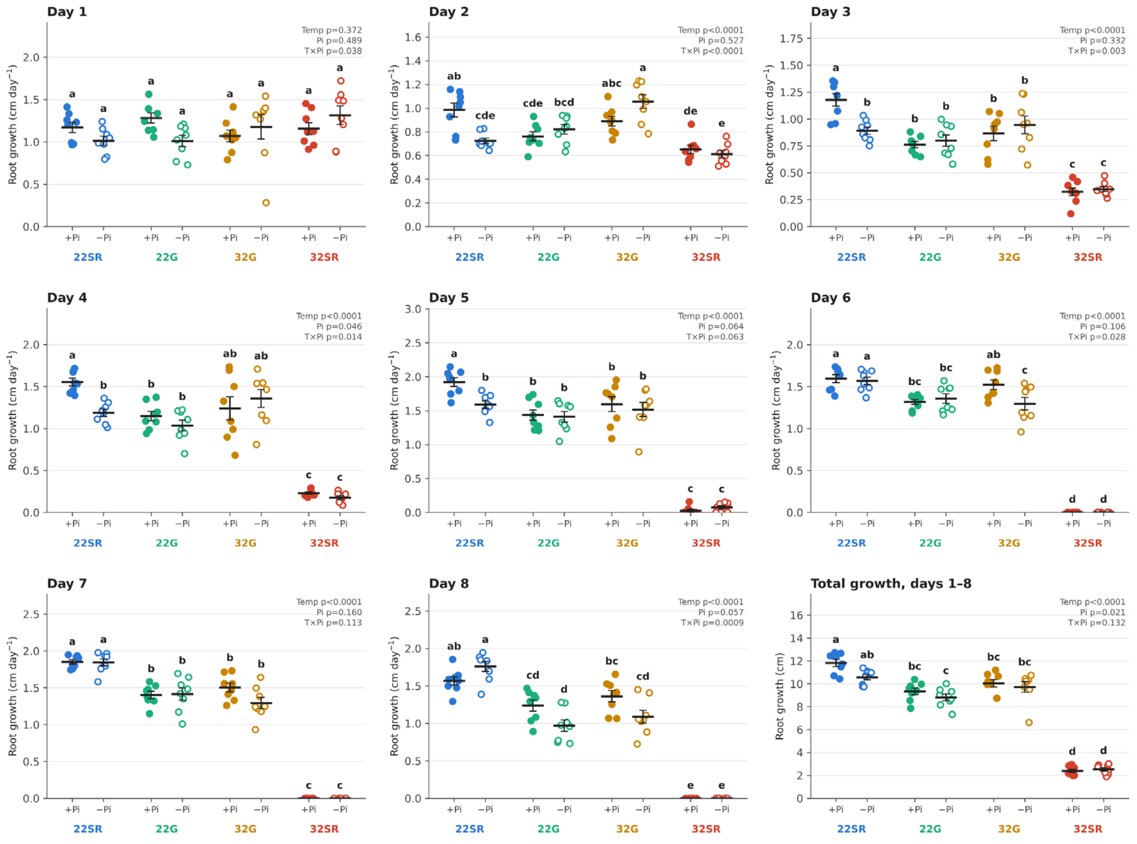


**Figure S2. Daily primary root growth of *Arabidopsis thaliana* seedlings under different temperature regimes and phosphate availability.** Four day-old seedlings grown under at 22SR-HPi or 22SR-LPi were transferred to homogeneous temperature of 22 °C (22SR) or homogeneous 32 °C (32SR). Seedlings grown under a 22 °C gradient (22G), were transferred to 22G or 32 °C TGRooZ (32G). Seedlings were grown at under high or low phosphate supply (HPi, 625 µM +Pi; LPi, 30 µM, -Pi). Panels Day 1–Day 8 show daily primary root growth (cm day$^{-1}$), where Day 1 is the growth after one days from the transference. Each dot is one seedling (n = 8). Horizontal black lines show the mean ± SEM. In each panel, the eight conditions were compared by one-way ANOVA followed by Tukey's HSD test; conditions that do not share a letter differ significantly ($p < 0.05$). The values in the upper right of each panel are the p-values of a two-way ANOVA testing the effects of temperature regime (Temp), phosphate availability (Pi) and their interaction (T×Pi). Under homogeneous 32 °C, root growth stopped completely from Day 6 onwards in both Pi treatments.

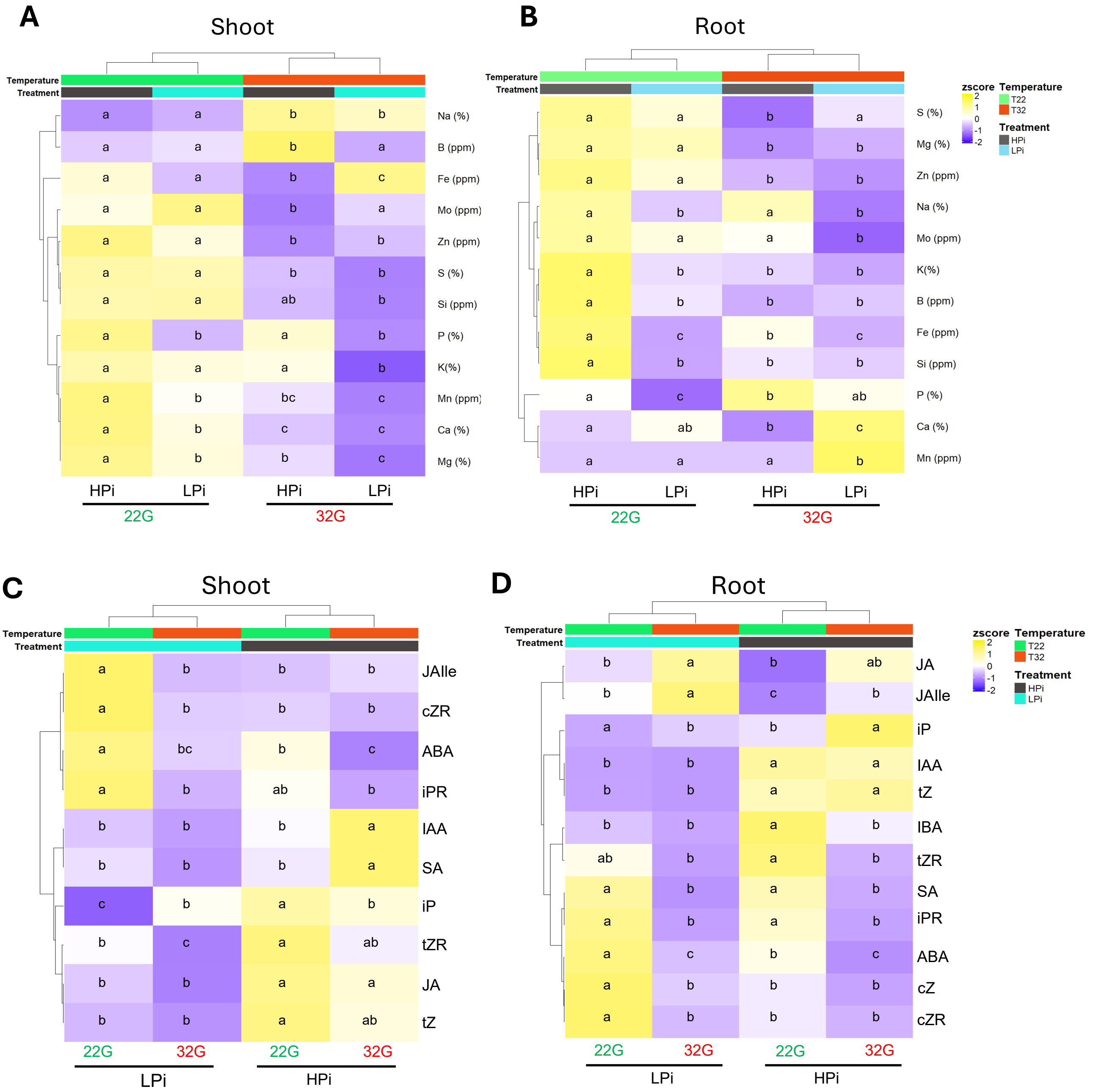


**Figure S3: Phosphate starvation and heat stress alter ion and hormone accumulation in roots and shoots.** (**A-B**) Heatmap of different ions in Arabidopsis shoot (**A**) or roots (**B**) grown in the TGRooZ at 22ºC or 32ºC (22G or 32G) with high or low phosphate (HPi - 625µM Pi; LPi 30 µM Pi) respectively. Values of different elements were presented as Z-scored of the percentage of the dried weight (%) or parts per million of dried weight (ppm) as indicated. (**C-D**) Heatmap of different hormone in Arabidopsis shoot (C) or roots (D) of seedlings grown in the TGRooZ at 22ºC or 32ºC (22G or 32G) with high or low phosphate (HPi - 625µM Pi) or LPi 30 µM Pi, respectively. Values of different hormones were presented as pmoles per gram of fresh weight. Different letter inside of the heatmap indicate the statistical differences between conditions by Tukey´s post hoc test, P < 0.05. The number of biological replicates were 4.

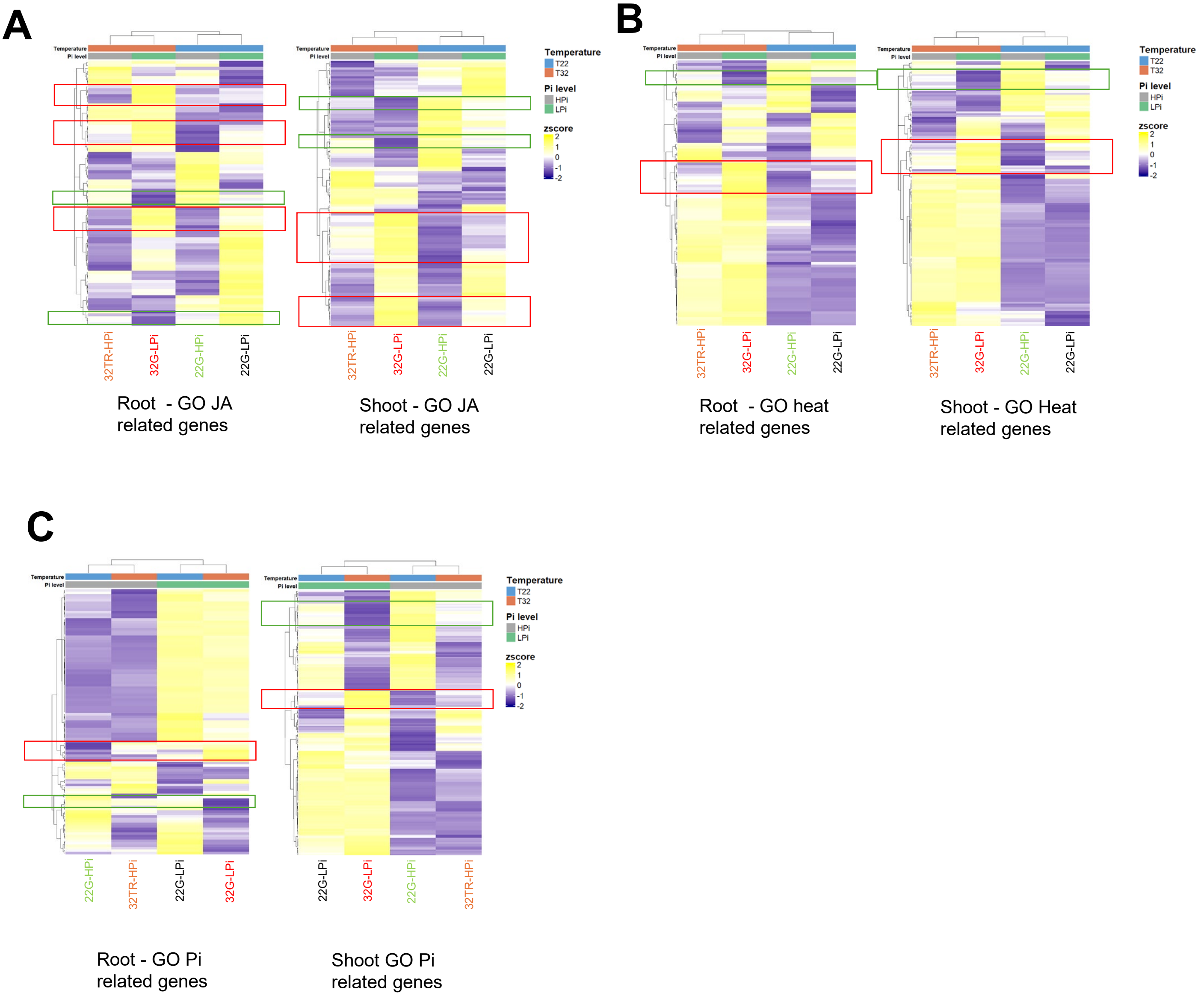


**Figure S4. Differential expression of genes associated with heat stress, phosphate signaling, and jasmonic acid pathways.** (A) Heat map showing the expression profiles of DEGs involved in jasmonic acid (JA)-related processes, including metabolism, homeostasis, transport, and signaling, across the different treatments (heat stress, phosphate deficiency, and combined stress). **(B-C)** Heat map showing the expression profiles of differentially expressed genes (DEGs) involved in heat stress responses (B) or phosphate-related processes (C), including metabolism, homeostasis, transport, and signaling, across the different treatments in root or shoot.

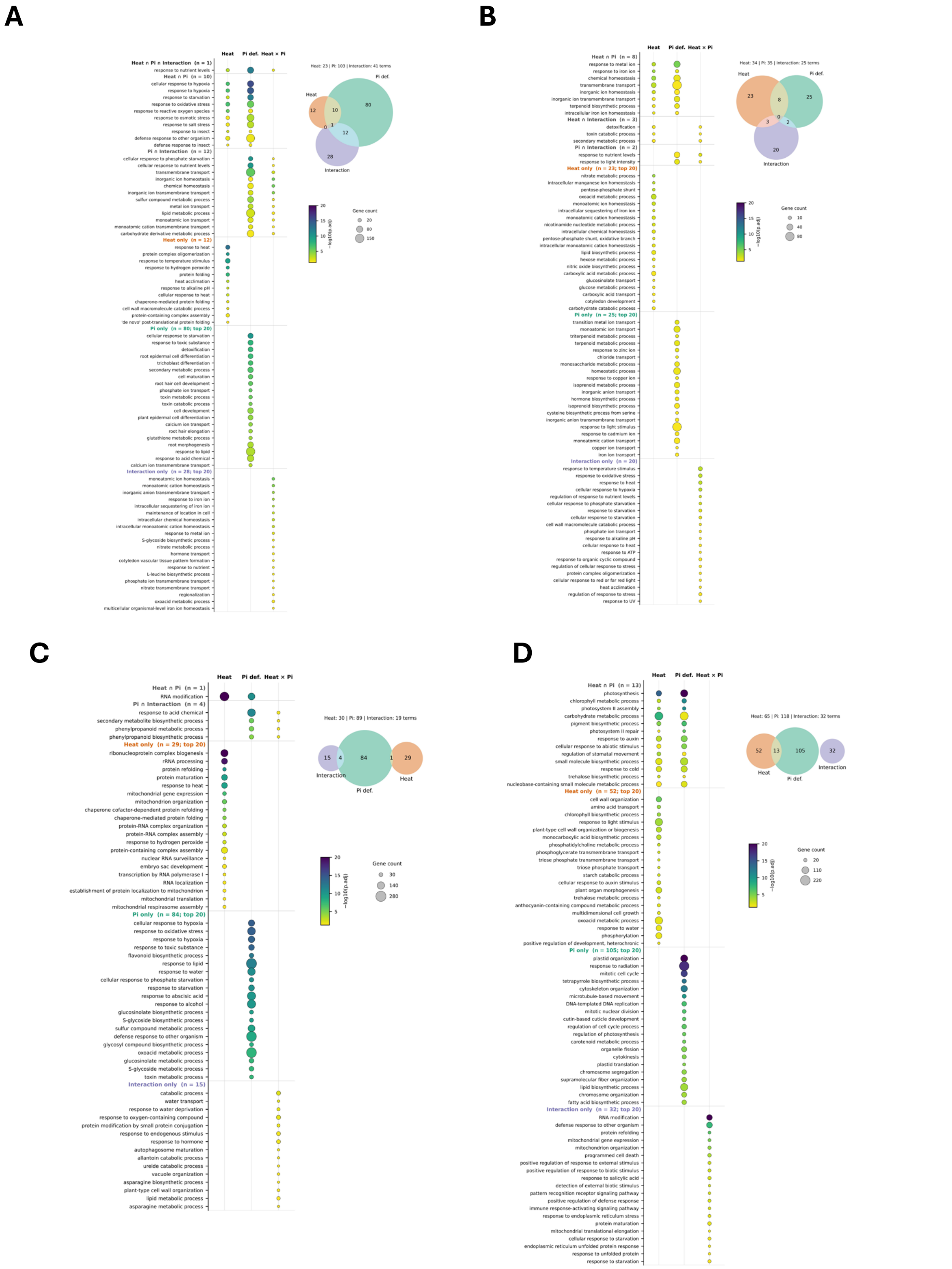


**Figure S5. Gene Ontology (GO) biological processes enriched among genes responsive to heat, Pi deficiency and their interaction in *Arabidopsis thaliana*.** GO enrichment of genes up- or down-regulated by heat, by Pi deficiency, and by the Heat × Pi interaction in roots and shoots. (A) Root, up-regulated genes. (B) Root, down-regulated genes. (C) Shoot, up-regulated genes. (D) Shoot, down-regulated genes. The Venn diagram in each panel shows the number of significantly enriched GO terms (adjusted p-value < 0.05) for heat, Pi deficiency and the interaction, and how many are shared between them. In the dot plots, GO terms are grouped by whether they are enriched in all three gene sets, in two of them (Heat ∩ Pi, Heat ∩ Interaction, Pi ∩ Interaction), or in only one (Heat only, Pi only, Interaction only). Within each group, terms are ordered by significance; for groups with more than 20 terms, only the 20 most significant are shown (the total number is indicated in brackets). Dot size is the number of genes annotated to each term, and dot color is the −log10 of the adjusted p-value (scale capped at 20). Interaction genes are those whose expression under combined heat and Pi deficiency differs from that expected from the additive effects of the two single stresses. GO enrichment was performed with clusterProfiler tool using the Arabidopsis genome as background, and p-values were adjusted by the Benjamini–Hochberg method.

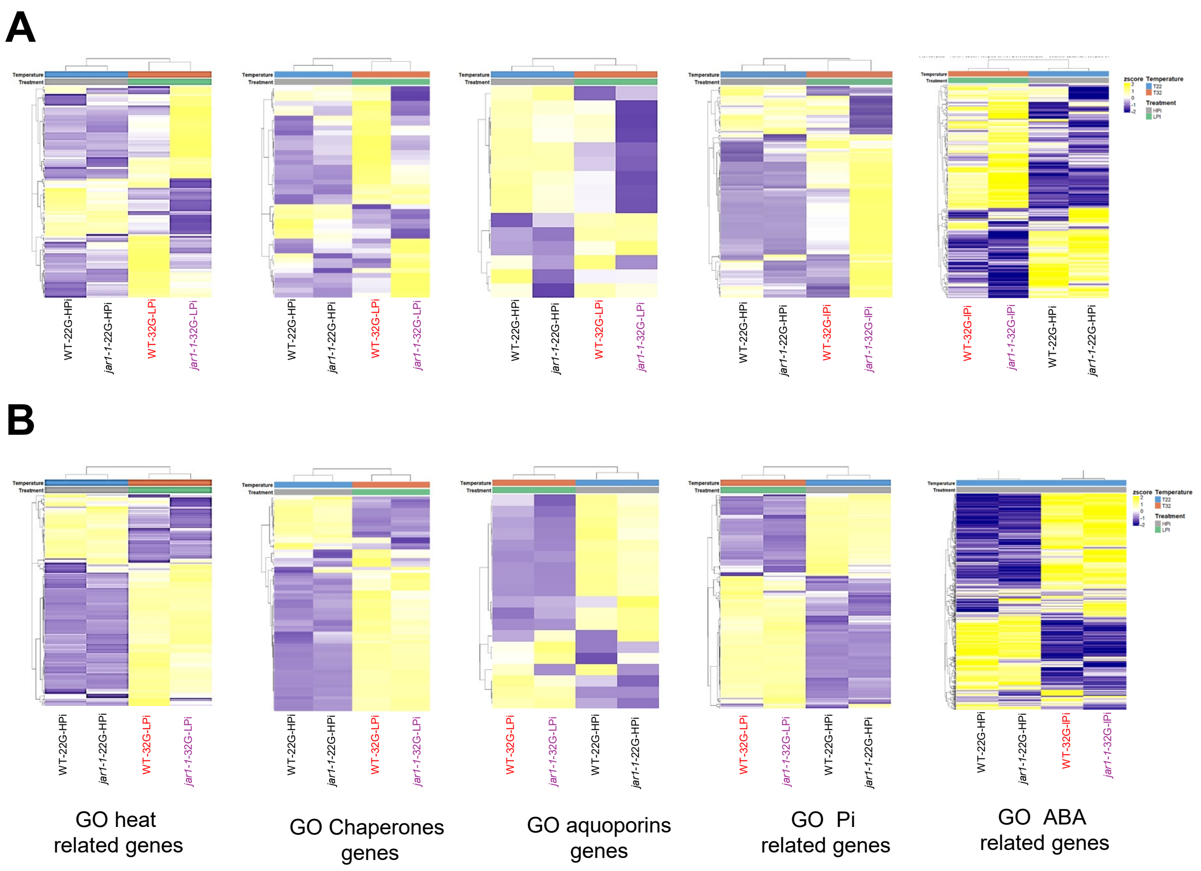


**Figure S6. Expression of heat-, chaperone-, aquaporin-, Pi- and ABA-related genes in WT and *jar1-1* plants under control and combined stress conditions.** Heat maps of transcripts identified by RNA-seq in **(A)** roots and **(B)** shoots of wild-type (WT) and *jar1-1* plants grown under control conditions (22 °C-TGRooZ, high Pi; 22G-HPi) or combined heat stress and Pi deficiency (32 °C-TGRooZ, low Pi; 32G-LPi). From left to right, panels show DEGs annotated with Gene Ontology (GO) terms related to heat stress responses, chaperones (including heat-responsive transcription factors), aquaporins, Pi homeostasis, and ABA signaling. Expression values are shown as row-scaled z-scores (yellow, higher expression; blue/purple, lower expression). Genes (rows) were grouped by hierarchical clustering. Colored bars above each heat map indicate growth temperature (22 °C or 32 °C) and Pi treatment (HPi or LPi). Sample labels are shown below each heat map: control samples in black, WT combined stress in red, and *jar1-1* combined stress in purple.
